# RfxCas13d Mediates Broad-Spectrum Suppression of Highly Pathogenic Avian Influenza

**DOI:** 10.64898/2026.03.18.712793

**Authors:** Sudip Dhakal, Aaron J. Smith, Emiliana Weiss, Md. Zohorul Islam, Lynn Nazareth, Terrence Lee, Tamara Gough, Kiran Krishnankutty Nair, Laurence Wilson, James W. Wynne, Kristie Jenkins, Arjun Challagulla

**Affiliations:** Health and Biosecurity, CSIRO, East Geelong, VIC, Australia; Agriculture and Food, CSIRO, Hobart, TAS, Australia; Health and Biosecurity, CSIRO, Black Mountain, ACT, Australia; Australian Animal Health Laboratory, CSIRO, East Geelong, VIC, Australia; Health and Biosecurity, CSIRO, Westmead, NSW, Australia

**Keywords:** RfxCas13d, highly pathogenic avian influenza virus, RNA targeting antivirals, multiplexed RNA targeting, Antiviral genome editing

## Abstract

Highly pathogenic avian influenza viruses (HPAIVs) continue to cause substantial disease in birds and mammals, with repeated H5N1 spillovers highlighting the need for broadly protective antiviral strategies. Here we develop a programmable RNA-targeting antiviral platform based on RfxCas13d and evaluate its activity in avian cells. Screening of five Cas13 orthologs in chicken DF1 fibroblasts revealed RfxCas13d as the most potent and well tolerated effector. Virus-specific CRISPR RNAs (crRNAs) targeting conserved regions of positive- and negative-sense influenza RNA were tested against A/WSN/033[H1N1] and multiple HPAIV isolates, including a member of clade 2.3.4.4b H5N1. Targeting positive-sense RNA conferred superior influenza inhibitory activity and further enhanced by multiplexed crRNA expression. These findings establish RfxCas13d as a versatile RNA-guided antiviral platform and provide a route for broad-spectrum influenza control through conserved RNA targeting.

## Introduction

Avian influenza viruses (AIVs) continue to cause enormous agricultural and public-health costs, fuelled by relentless evolution and repeated cross-species transmission. AIVs are classified as low pathogenic (LPAIV) or highly pathogenic (HPAIV) based on their pathogenicity. LPAIVs circulate endemically in wild waterfowl and typically cause asymptomatic infections. However, upon introduction into poultry, certain H5 and H7 strains can mutate to acquire multibasic cleavage sites in the hemagglutinin gene, converting into highly pathogenic forms and enabling systemic infection ^1,2^. The rapid viral evolutionary dynamics, driven by the error-prone RNA polymerase, frequent genetic reassortment, and strong immune selection, facilitate these evolutionary shifts and the emergence of novel HPAI strains with increased virulence and expanded host range ^3^.

In poultry, the first HPAIV outbreak was recorded in Scotland in 1959 with the “Smith” strain (A/Chicken/Scotland/59[H5N1]) ^4^. As early as 1997 in Hong Kong, HPAI H5N1 has been detected to cause human infections resulting on human fatalities ^5^. Between 2016 and 2020, the evolution of H5N1 viruses from clade 2.3.4.4b has noticeably shifted the range of host leading to mass mortalities of several avian species ^6–8^. Since then, incursions of the members of clade 2.3.4.4b have been reported in geographically isolated regions and across an expanded host range to different species of animals, including cattle and humans ^9–12^. The alarming spread of the HPAIV to global and rapid widespread multi-year outbreaks across various animal species increases concern for the global poultry production and food security ^13,14^. Even more concerning is its enzootic occurrence in wild migratory birds which has resulted in the spread of the HPAI virus in exotic regions of the world such as Antarctica ^15–17^. Multiple mitigation strategies, including enhanced surveillance, strengthened biosecurity, antiviral and vaccine development, closure of live bird markets, and culling of infected flocks, have been implemented to control viral spread ^14^. Despite the multilayered control measures being applied to prevent the HPAIV threats, the current disease preparedness strategies are falling short with increased reports of incursions of the HPAIV outbreaks ^18^. This persistent threat reinforces the urgent need for innovative interventions. One promising avenue is the genetic engineering of avian influenza–resilient poultry, capable of restricting viral replication in chickens and thereby mitigating zoonotic transmission risk.

Originally evolved as bacterial defence system against invading phage and plasmids, Clustered regularly interspaced palindromic repeats (CRISPR)/Cas systems have been repurposed as a powerful genetic tool with broad applications, including antiviral strategies ^19–21^. The first and most widely used CRISPR/Cas9 system, is a DNA targeting effector which causes double stranded breaks on DNA strands based on the guide RNA (gRNA)-mediated sequence complementarity. In previous studies, CRISPR/Cas9 has been used as antiviral approach against viruses, including human immunodeficiency virus (HIV) and hepatitis B virus. In these studies, viral inhibition was achieved either by excising and editing viral genes or by knocking out important host genes that has critical role in the viral replication ^22–25^. In poultry, CRISPR/Cas9 has been deployed in transgenic models to confer resistance against Marek’s disease virus, demonstrating the feasibility of direct targeting of viral genome as an effective antiviral strategy ^26^. However, CRISPR/Cas9-based targeting is inherently constrained to target DNA viruses or viruses with DNA intermediates during replication cycle. In the context of RNA viruses such as influenza A virus, viral replication is carried out exclusively by the RNA genome, rendering CRISPR/Cas9 ineffective for direct targeting of influenza virus genome. To this end, RNA-targeting systems such as CRISPR/Cas13 system, with their ability to recognise and cleave RNA in a sequence specific manner, offer a powerful alternative tool, including influenza virus.

Cas13 effectors, part of the Class II Type VI CRISPR-Cas system, consist of a single, large, multi-domain protein and function as a programmable, RNA-guided RNA-targeting system ^27^. Upon target RNA recognition, the Cas13 effector undergoes conformational changes to its structure activating its *cis*-cleavage action in the sequence complementary to the RNA guide or CRISPR RNA (crRNA). In addition, the activated effector also exhibits *trans*-cleavage activity, leading to degradation of non-specific RNA molecules present in its close vicinity, a phenomenon termed as collateral activity. While several variants of the Cas13 effectors have been identified in a recent metagenomic discoveries across bacterial community ^28^, their mechanistic features, target specificity and collateral activity remain poorly understood. As RNA-targeting effectors emerge as promising candidates for next-generation antivirals, understanding how to exploit their ability to directly inhibit viruses by targeting viral RNA has become increasingly imperative ^29^.

Although CRSIPR/Cas13 systems have been shown to reduce influenza viruses in human cell lines and mouse models ^30–32^, their application in transgenic poultry and efficacy against highly pathogenic avian influenza viruses have not yet been investigated. Since, the HPAI viruses evolve rapidly and known to have higher mortality and evolutionary rates, evaluating the efficacy of CRISPR/Cas13 system against the emerging HPAI viruses provided rationale to this study. In this study, we evaluated the antiviral potential of transgenic chicken cells modified to express RfxCas13d against HPAI viruses. Our results indicate that RfxCas13d can limit virus replication *in vitro*. The extent of reduction varied depending on the target selection and the virus subtype used. Viral inhibition was further enhanced by multiplexing crRNAs. These findings support the potential of RfxCas13d as a tool to mitigate HPAI infection.

## Results

### 1. Systematic Comparison of Target-Specific and Collateral Activities of Cas13 Effectors Using a Three-Colour Fluorescence Assay in chicken cells

A key consideration in the application of Cas13-based antiviral strategies is its collateral RNase activity, whereby target recognition can trigger nonspecific cleavage of surrounding RNA. To systematically evaluate the target-specific and collateral activities of five Cas13 effectors (LwaCas13a, PspCas13b, RfxCas13d, Cas13e3, and Cas13x(HF)), three-colour fluorescence reporter assay (Vector maps provided in Fig. S1, S2) was developed in chicken embryonic fibroblast (DF1) cells. Schematic of methods employed for this screening is shown in Fig. S3a. Transient co-transfection of Cas13 expression vectors (selection marker: miRFP670nano, red fluorescence), crRNA vector (selection marker: mTagBFP2, blue fluorescence) targeting green fluorescent protein (GFP) messenger RNA, and a GFP reporter plasmid (green fluorescence) enabled simultaneous assessment of Cas13 expression, target specific knockdown, and collateral activity. Expression of miRFP670nano served as an internal control for transfection efficiency, which inversely correlated with vector size, with larger Cas13 vectors yielding lower red fluorescence intensity and fewer miRFP670nano-positive cells (Fig. 1a, b, Fig. S3b). Target-specific activity was assessed by GFP knockdown, visualized by fluorescence microscopy (Fig. 1c) and quantified via flow cytometry (Fig. 1d, e). RfxCas13d demonstrated the highest target specific knockdown efficiency, reducing GFP fluorescence by >90%, followed by Cas13e3 (∼30%) and LwaCas13a (∼10%), respectively (Fig. 1d, e). In contrast,

**Fig. 1:**
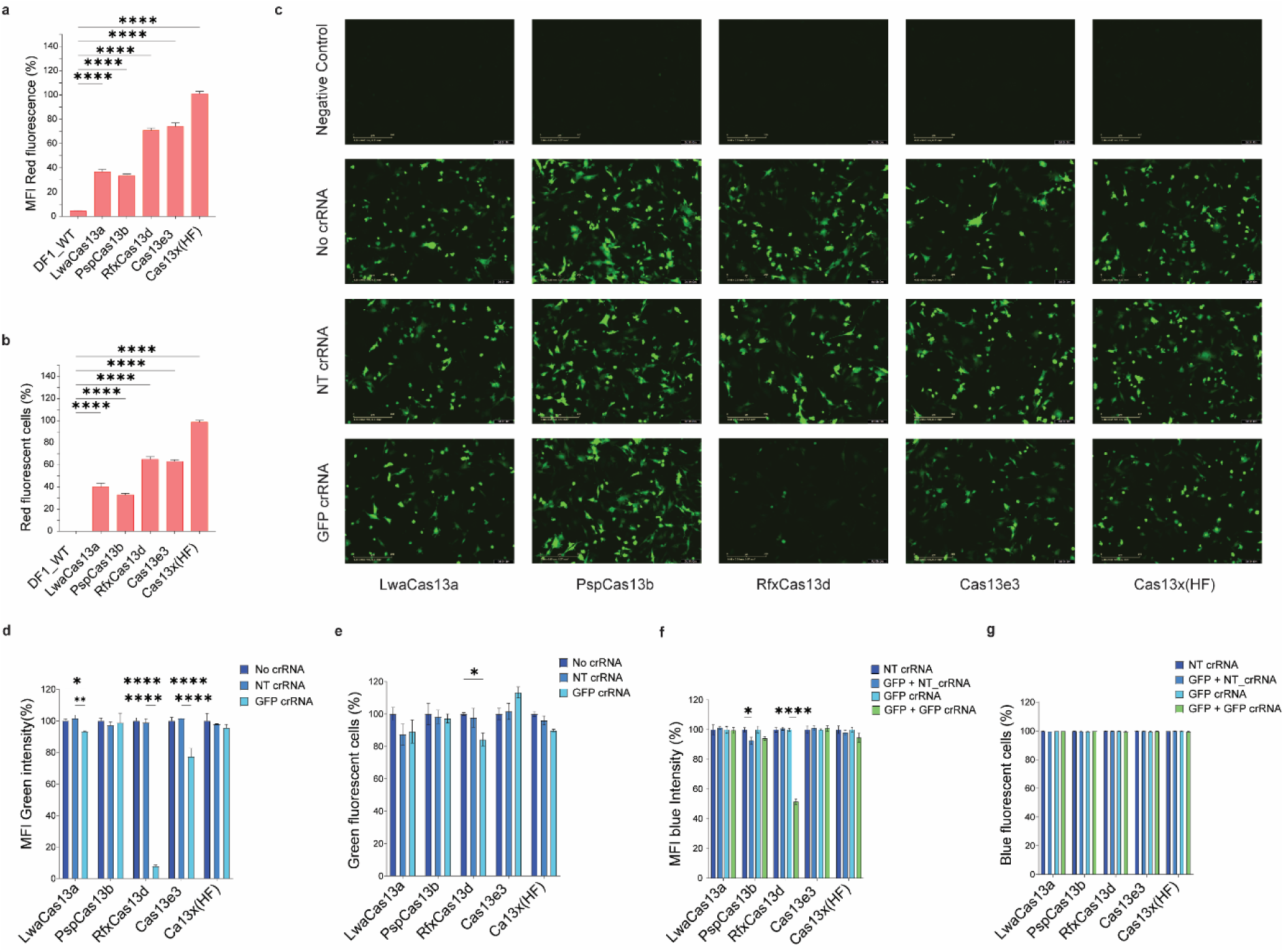
Functional screening of CRISPR–Cas13 systems for target-specific and collateral activity in chicken DF1 cells. **a, b** Flow cytometry-based quantification of Cas13 vector expression across variants (LwaCas13a, PspCas13b, RfxCas13d, Cas13e3, Cas13x(HF)) in DF1 cells by measuring mean fluorescence intensity (MFI) of miRFP670nano (a) and percentage of miRFP670nano-positive cells (b). **c** Representative green fluorescence micrographs of DF1 cells co-transfected with Cas13 variants, either target-specific crRNA (GFP crRNA) or non-targeting crRNA (NT crRNA) or without crRNA (No crRNA), and a GFP reporter plasmid. **d, e** Evaluation of GFP knockdown efficiency via MFI (d) and number of GFP-positive cells (e). **f, g** Assessment of collateral activity through quantification of mean fluorescence intensity (MFI) (f) and mTagBFP2-positive cells (g). Data for miRFP670nano expression were analysed using one-way ANOVA with non-transfected DF1 cells as control. All other datasets were analysed using two-way ANOVA with GFP + No crRNA and GFP + NT crRNA conditions as reference. Statistical significance is indicated as follows: *p < 0.05; **p < 0.01; ***p < 0.001; ****p < 0.0001.

PspCas13b and Cas13x(HF) showed negligible on-target activity under the same conditions. Quantification of cellular ATP (as a measure of cell viability) revealed a modest reduction in viability only in Cas13e3 transfected cells expressing target-specific crRNA in the presence of GFP (target), whereas all other treatments-maintained viability comparable to controls (Fig. S3c). Collateral activity was evaluated by monitoring mTagBFP2 fluorescence, expressed from the crRNA vectors. RfxCas13d induced a significant reduction (∼50%) in blue fluorescence, indicating a moderate collateral degradation of non-target transcripts (Fig. 1f, g; Fig. S3d). The remaining Cas13 effectors did not show any measurable reduction of mTagBFP2 fluorescence. In summary, RfxCas13d is the most effective RNA-targeting Cas13 system among the five Cas13 variants tested in chicken cells. Although the potent target-specific knockdown of RfxCas13d was accompanied by moderate collateral activity against non-target transcripts, the bystander effect did not compromise cell viability.

### 2. Selective targeting of nuclear viral RNA species by RfxCas13d reveals differential antiviral efficacy

Given that influenza virus replication involves the generation of both positive- and negative-sense viral RNA intermediates within the nucleus of infected cells, we investigated the antiviral activity of RfxCas13d by designing crRNAs against both classes of viral RNA intermediates. For targeting the positive-sense viral RNA, nine crRNAs were generated against three viral segments: PB2 (PB2_M1–M3), PB1 (PB1_M4–M6), and NP (NP_M7–M9), and screened using a transient transfection assay (crRNA design strategy refer to Fig. S4). The screening workflow for antiviral activity by transient crRNA transfection followed by infection and viral M-gene evaluation is illustrated in Fig. S5a. Following this initial screening, the best performing crRNA candidates were chosen to generate stable RfxCas13d-crRNA targeting positive sense RNA, along with the negative sense RNA targeting crRNAs. Schematic of the methods employed to test the crRNAs in stable RfxCas13d cell lines are shown in Fig. S5b. Transient transfection of individual crRNAs into stable RfxCas13d-expressing cells, followed by A/WSN/033[H1N1] infection and quantitative RT-PCR, revealed that PB1_M4 and NP_M7 significantly reduced viral RNA levels by approximately 1-fold compared to non-targeting (NT crRNA) (Fig. 2a). Based on these findings, PB1_M4, PB1_M6 and NP_M7 were selected for stable expression in RfxCas13d cell lines and tested further. Upon infection of stable RfxCas13d-crRNA cells with A/WSN/033[H1N1], PB1_M4 and NP_M7 stable cells exhibited robust antiviral activity (Fig. 2b–d). Specifically, NP_M7 reduced viral titres by ∼2.3 log units, while crRNA_M4 achieved a ∼1.4 log unit reduction at 24 hours post-infection (h.p.i). In contrast, crRNA_M6 did not confer measurable antiviral activity (Fig. 2b-d). Corresponding cytopathic effects observed at 72 h.p.i in these cell lines are shown in Fig. S5c.

**Fig. 2:**
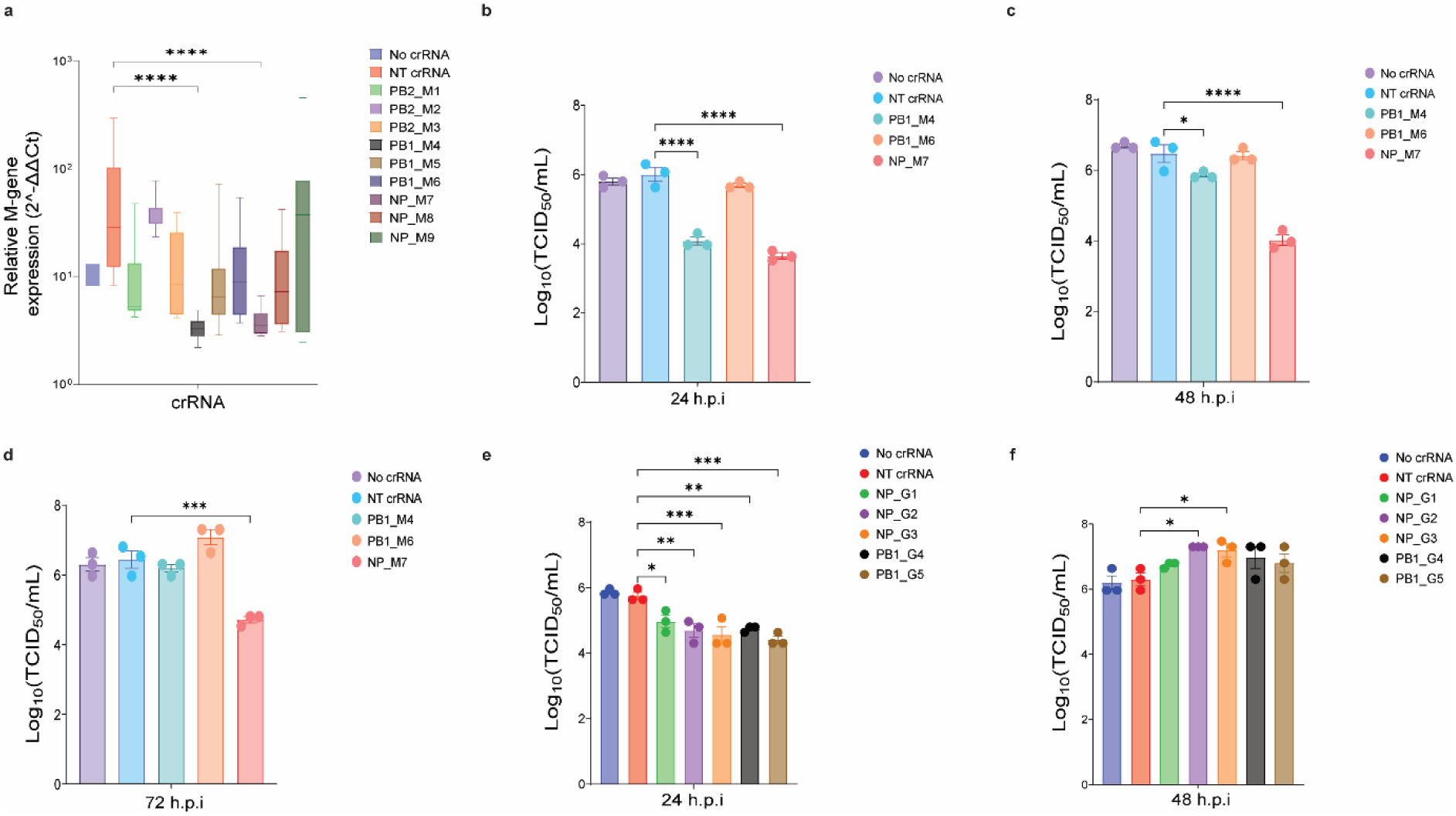
Comparative antiviral efficacy of crRNAs targeting positive and negative sense influenza viral RNAs in RfxCas13d-expressing cell lines. **a** Relative fold change of M-gene (2^-ΔΔCt^) in RNA extracted from RfxCas13d cell lines transiently transfected with crRNAs targeting positive sense RNA (PB2_M1-M3, PB1_M4-M6 and NP_M7-9) and infected with A/WSN/033[H1N1] at 24 h post-transfection. **b-d** Viral titres determined by TCID₅₀ assay in MDCK cells from supernatants of infected DF1 cell lines stably expressing RfxCas13d (No crRNA), non-targeting crRNA (NT crRNA), PB1_M4, PB1_M6, and NP_M7 at 24 h.p.i (b), 48 h.p.i (c), and 72 h.p.i (d). **e, f** Viral titres in DF1 cell lines stably expressing No crRNA, NT crRNA, or crRNAs targeting negative sense RNA (NP_G1–G3 and PB1_G4-G5) at 24 h.p.i (e) and 48 h.p.i (f). Viral titres were calculated using the Reed–Muench method and analysed by one-way ANOVA, with comparisons made to NT crRNA controls. Statistical significance is indicated as follows: *p < 0.05; **p < 0.01; ***p < 0.001; ****p < 0.0001.

To evaluate the targeting of negative-sense genomic RNA, five additional crRNAs (NP_G1-G3 and PB1_G4-G5) were designed against NP and PB1 segments. Due to variability observed in transient transfection assays, these constructs were tested directly using stable RfxCas13d-crRNA cell lines. Following A/WSN/033[H1N1] infection, all five cell lines demonstrated antiviral activity, with reductions in viral titres ranging from 0.7 to 1.3 log units at 24 h.p.i (Fig. 2e). However, by 48 h.p.i, viral titres progressively increased across all cell lines, reaching levels comparable to or exceeding non-targeting (NT crRNA) controls (Fig. 2f). Corresponding cytopathic effects shown by these negative sense RNA targeting RfxCas13d-crRNA cells are presented in Fig. S5d. Taken together, these results demonstrate that the RfxCas13d confers effective antiviral activity against influenza virus A/WSN/033[H1N1] with both positive-sense and negative-sense viral RNA targeting crRNAs. Notably, targeting the positive-sense RNA species within the nuclear compartment yielded a more robust antiviral effect than targeting the negative-sense (genomic) RNA, highlighting a potential mechanistic advantage in antiviral strategy design.

### 3. crRNAs targeting positive-sense RNA mediate potent and subtype-specific RfxCas13d antiviral activity

To evaluate the antiviral efficacy of RfxCas13d against highly pathogenic avian influenza viruses (HPAIVs), we adapted or reused previously characterized crRNAs based on sequence similarity. Two crRNAs targeting positive-sense RNA (PB1_M4 and NP_M7) and three targeting negative-sense RNA (NP_G1, PB1_G4, and PB1_G5) were selected to target conserved regions of H5 (A/Chicken/Vietnam/08/2004[H5N1]) and H7 (A/Chicken/Lethbridge/09/2020[H7N7]) subtypes. Modified crRNAs (NP_M10 and PB1_M11), derived from NP_M7 and PB1_M4 respectively, were designed to improve compatibility with HPAIV sequences. Schematic of methods employed to study the antiviral effects of selected RfxCas13d-crRNA cells is shown in Fig 3a.

**Fig. 3:**
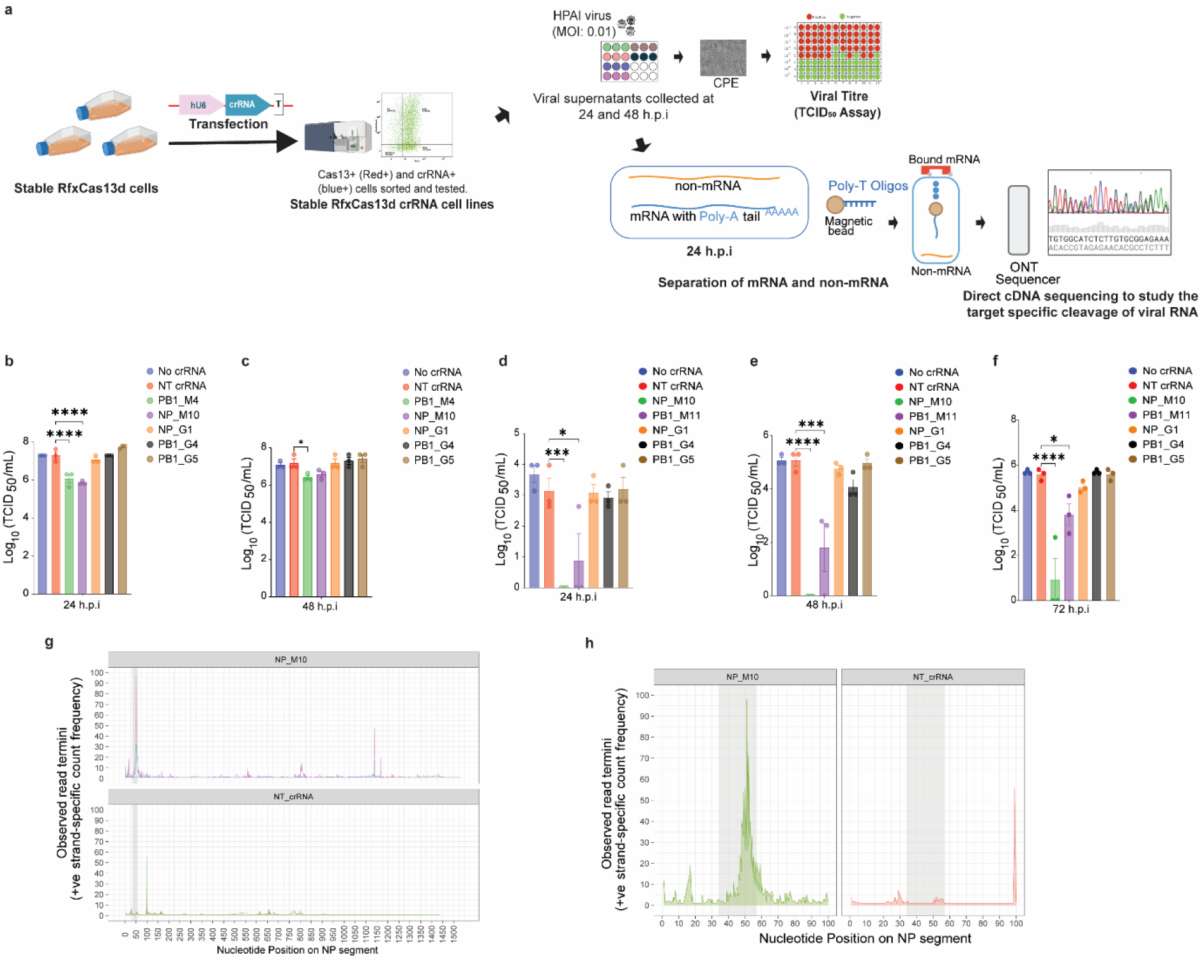
RfxCas13d restricts avian influenza virus replication and induces guide-dependent viral RNA termination. **a** Schematic of method used to evaluate the antiviral activity of RfxCas13d cell lines against HPAI viruses of H5 and H7 subtypes and to analyse guide-dependent viral RNA cleavage by long-read sequencing. **b-c** RfxCas13d cells expressing No crRNA, non-targeting crRNA (NT crRNA), or crRNAs targeting influenza virus (PB1_M4, NP_M10 and PB1_M11) were infected with A/Chicken/Vietnam/08/2004[H5N1] and viral titres determined by TCID50 assay in MDCK cells from supernatants at (b) 24 h.p.i and (c) 48 h.p.i. **d-f** Antiviral activity against A/Chicken/Lethbridge/9/2020[H7N7] was assessed by TCID50 assay in MDCK cells from viral supernatants at (d) 24 h.p.i, (e) 48 h.p.i, and (f) 72 h.p.i. Viral titres were calculated using the Reed–Muench method. Statistical significance is indicated as p < 0.05 (*), p < 0.01 (**), p < 0.001 (***), and p < 0.0001 (****). **g** Genome-wide alignment of NP positive strand cDNA read termini from A/Chicken/Vietnam/08/2004[H5N1] infected RfxCas13d cells expressing either NT crRNA or NP_M10 at 24 h.p.i. **h** Pile-up of cDNA read termini within the NP_M10 crRNA spacer-matched region (grey shading), indicating guide-dependent enrichment of transcript termination events.

Infection of stable RfxCas13d-crRNA cell lines with HPAIVs revealed viral subtype-dependent antiviral activity. At 24 h.p.i, NP_M10-expressing cells showed a 1.4 log unit reduction in viral titre of A/Chicken/Vietnam/08/2004[H5N1], whereas PB1_M4-expressing cells showed a 1.3 log unit reduction (Fig. 3b). By 48 h.p.i, only PB1_M4 maintained significant suppression of viral replication (Fig. 3c). In contrast, infection of NP_M10 expressing cells with A/Chicken/Lethbridge/09/2020[H7N7] showed no detectable viral load (>5 log unit reduction), sustained through 48 h.p.i (Fig. 3d, e). However, PB1_M11-expressing failed to exhibit antiviral activity across all time points (24, 48, and 72 h.p.i), highlighting the enhanced antiviral activity of the NP_M10 crRNA against the H7N7 strain (Fig. 3c–e). Notably, cell lines expressing negative-sense RNA-targeting crRNAs (NP_G1, PB1_G4, PB1_G5) failed to elicit measurable antiviral effects against HPAIV subtypes. Corresponding cytopathic effects of the RfxCas13d cells infected with HPAIVs A/Chicken/Vietnam/08/2004[H5N1] and A/Chicken/Lethbridge/09/2020[H7N7] are shown in Fig. S6 and Fig. S7, respectively.

To examine the cis-cleavage activity and cleavage pattern of RfxCas13d on viral RNA, total RNA was isolated from A/Chicken/Vietnam/08/2004[H5N1] infected cells expressing RfxCas13d together with either NP_M10 crRNA or NT crRNA. Messenger RNA was enriched from total RNA using Dynabead-based purification and analysed by Oxford Nanopore long-read direct cDNA sequencing (Fig. 3a). Viral genome-wide mapping of cDNA reads revealed a single region of increased read pile-up along the NP transcript in the NT crRNA, consistent with a background site of coverage enrichment independent of guide targeting. In contrast, cells expressing the NP_M10 displayed multiple regions of increased read pile-up across the NP transcript (Fig. 3g). Notably, the most prominent accumulation of reads was centred on the NP_M10 guide target region, where a high density of reads exhibited 5′ termini terminating within the spacer-matched sequence and frequent 5′ soft-clipping. The highest concentration of read termini mapped to positions 47–55 of the NP genome, corresponding to a 13–21 bp window within the crRNA spacer target region, whereas enrichment at this site was minimal in the NT crRNA condition (Fig. 3h). Together, these observations indicate that guide-dependent targeting by RfxCas13d is associated with reproducible and localized RNA termination events within the crRNA-matched region, consistent with a cis-cleavage mechanism in which RfxCas13d generates defined cleavage sites on the target RNA.

To define the mechanistic basis underlying the differential antiviral efficacy of distinct crRNAs, we compared early viral RNA kinetics at 6 and 24 h.p.i in RfxCas13d-expressing cells harbouring either the positive-sense targeting PB1_M4 crRNA or the negative-sense targeting PB1_G5 crRNA following infection with A/Chicken/Vietnam/08/2004[H5N1] (schematic in Fig. 4a). Quantitative RT–PCR analysis at 6 h.p.i revealed that PB1_M4 significantly reduced all three viral RNA intermediates, genomic (vRNA), complementary (cRNA) and messenger RNA (mRNA), whereas PB1_G5 selectively reduced vRNA with minimal effects on cRNA and mRNA (Fig. 4b–d).

**Fig. 4.**
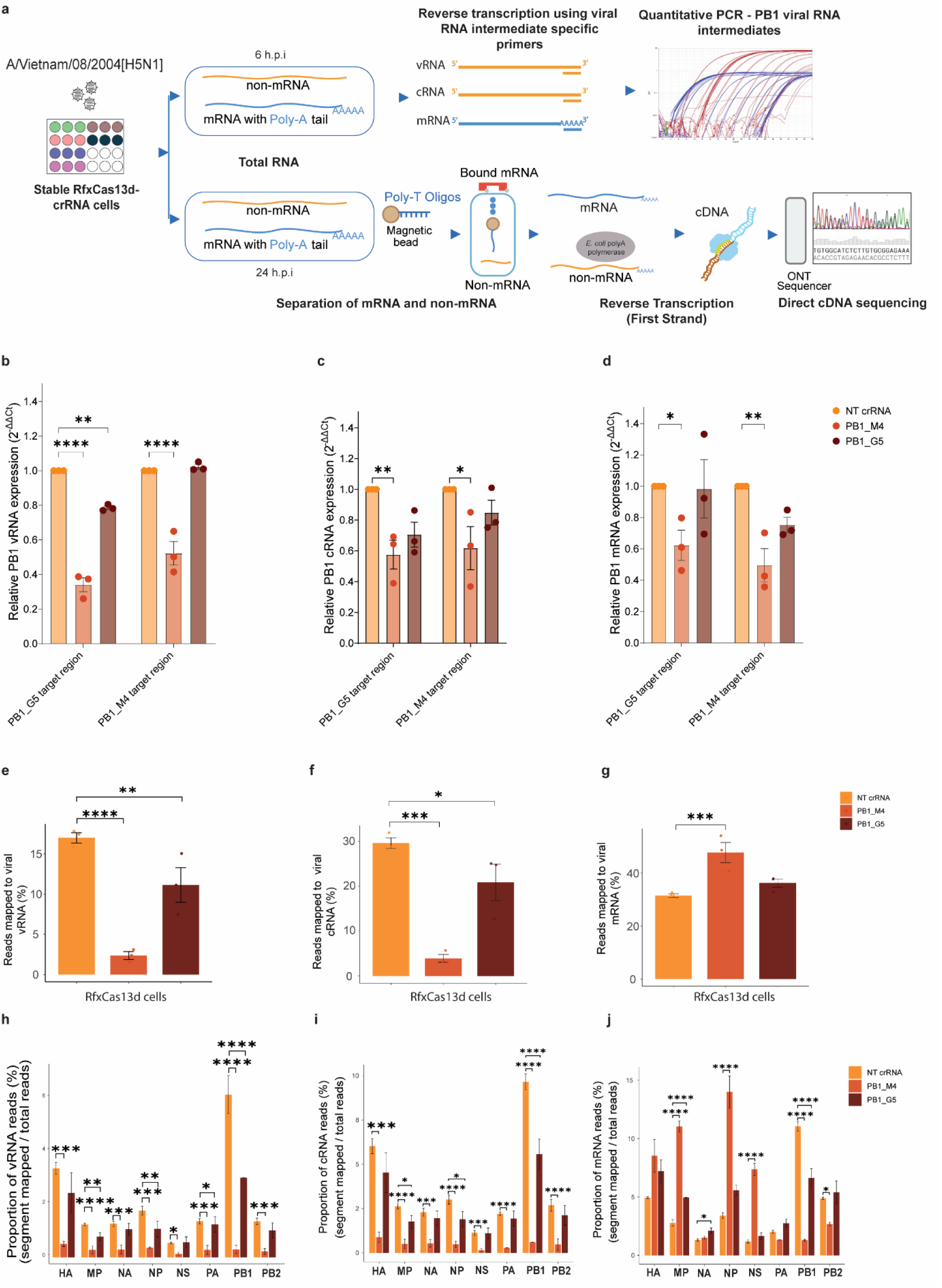
Strand orientation of crRNAs determines RfxCas13d antiviral activity. **a** Experimental workflow used to compare the antiviral efficacy of positive-sense– and negative-sense–targeting crRNAs in RfxCas13d-expressing cells. **b–d** Effects of crRNA targeting on viral RNA intermediates were assessed by quantitative RT–PCR analysis of the PB1 segment in RfxCas13d cell lines stably expressing PB1_M4 or PB1_G5 crRNAs and compared with non-targeting (NT) crRNA controls. Relative RNA expression levels (2^−ΔΔCt^) are shown for viral genomic RNA (vRNA) (b), complementary RNA (cRNA) (c), and messenger RNA (mRNA) (d), measured using TaqMan probes targeting the PB1_M4 and PB1_G5 target regions. Data represent the mean ± standard error from mean from three independent biological replicates (n = 3). Statistical significance was determined by one-way ANOVA relative to RfxCas13d NT crRNA cell lines. **e–j** Global and segment-specific distributions of viral RNA read obtained by Oxford Nanopore long-read direct cDNA sequencing. Shown are the proportions of mapped vRNA (e), cRNA (f) and mRNA (g) reads in RfxCas13d cells expressing NT, PB1_M4 or PB1_G5 crRNAs, together with segment-wise analyses of vRNA (h), cRNA (i), and mRNA (j) abundance across individual viral segments. Sequencing data are representative of three independent biological replicates (n = 3). Statistical significance was determined by paired t-test relative to RfxCas13d NT crRNA cell lines. Statistical significance is indicated as p < 0.05 (*), p < 0.01 (**), p < 0.001 (***), and p < 0.0001 (****).

To independently validate these observations, total RNA isolated at 24 h.p.i was subjected to Oxford Nanopore long-read sequencing using a direct cDNA approach. Messenger RNA was enriched using Dynabeads, while non-mRNA fractions were purified and poly(A)-tailed prior to library preparation. mRNA and non-mRNA libraries were prepared using identical workflows, barcoded separately, sequenced independently, and analysed to quantify viral RNA intermediates. Consistent with the qRT–PCR data, genome-wide analysis demonstrated reduced overall levels of vRNA and cRNA in both PB1_M4 and PB1_G5-expressing cells relative to NT crRNA controls (Fig. 4e, f). The mRNA levels in both the cells either increased or remain same relative to the NT crRNA controls (Fig. 4g). Interestingly, across nearly all samples and conditions, positive sense cRNA reads mapping to each viral genome segments were approximately two-fold higher than their vRNA counterparts, suggesting a greater abundance of positive-sense RNA within infected cells (Fig. 4e, f, h, i).

Segment-resolved mapping revealed a more pronounced and uniform reduction of vRNA and cRNA across all viral segments in PB1_M4-expressing cells (Fig. 4h, i). In contrast, PB1_G5 expression resulted in significant reductions in vRNA levels for the MP, NP, PA and PB1 segments and in cRNA levels for MP, NP and PB1 only (Fig. 4h, i). Notably, the magnitude of vRNA reduction was consistently greater in PB1_M4-expressing cells than in PB1_G5-expressing cells. Conversely, analysis of mRNA abundance revealed a complex segment-specific response, with increased levels of MP, NP and NS mRNAs in PB1_M4 cells, alongside a marked reduction in PB1 and PB2 mRNAs (Fig. 4j).

Collectively, these findings indicate that positive-sense targeting crRNAs can simultaneously impact multiple viral RNA intermediates, thereby conferring enhanced antiviral activity compared with negative-sense targeting. Furthermore, the differential effects observed across viral RNA species and segments suggest that the stage of the viral replication cycle and RNA strand accessibility critically shape the outcome of RfxCas13d-mediated antiviral suppression.

### 4. Ribozyme-based crRNA arrays enable consistent multiplexed guide activity in RfxCas13d systems

To determine whether multiplexed crRNA expression enhances RfxCas13d-mediated inhibition of influenza A virus, we compared two crRNA array architectures: (i) a self-cleaving ribozyme (Hammerhead and Hepatitis Delta Virus ribozymes), and (ii) a chicken glycyl tRNA sequences for crRNA processing (Fig. 5a). Each array construct contained three crRNAs; two non-targeting controls and one GFP-targeting guide, cloned into a standard expression vector. These were assessed alongside single-guide GFP_crRNA and non-targeting controls for target-specific activity (Fig. 5a).

**Fig. 5:**
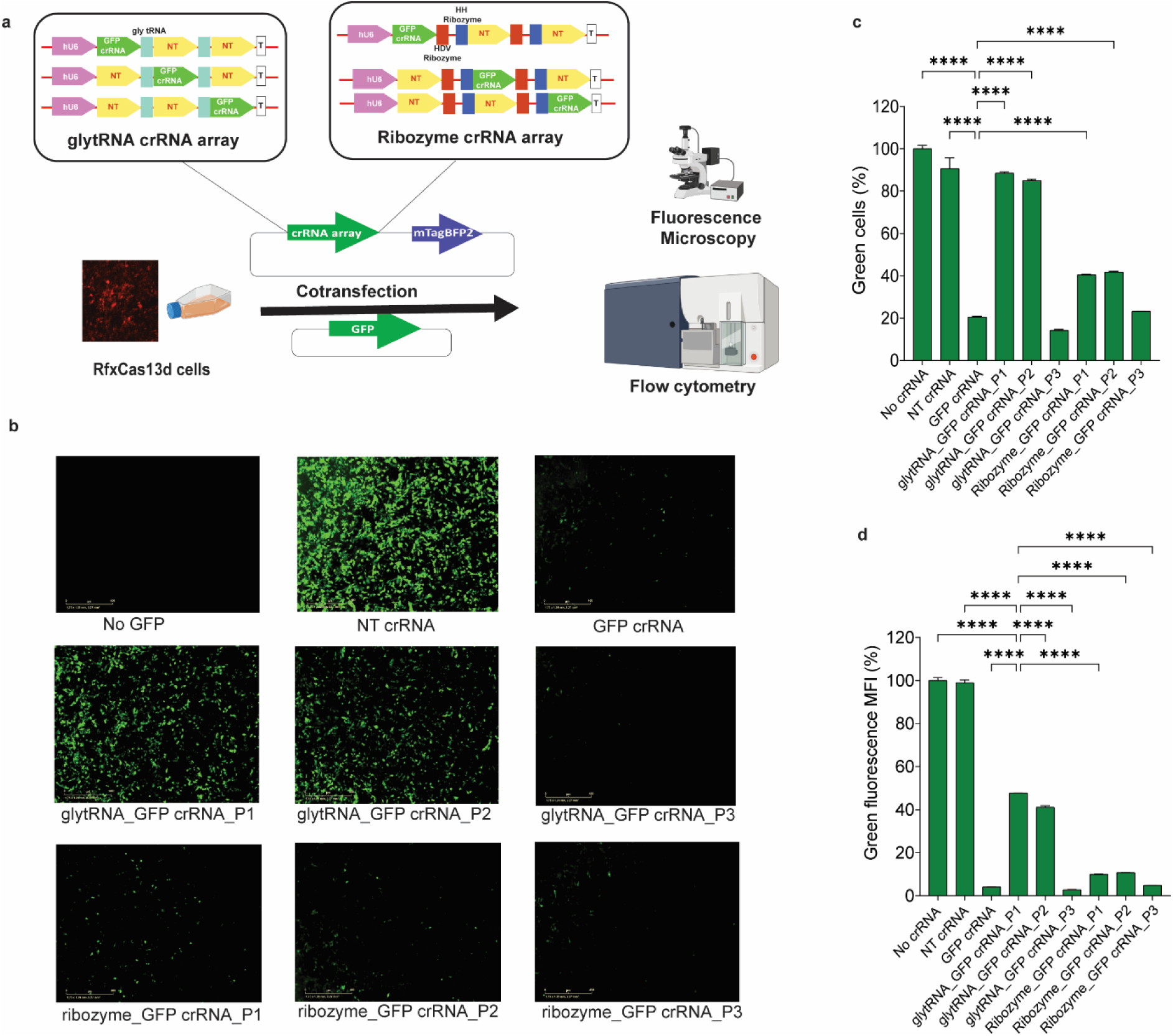
Evaluation of crRNA multiplexing strategies for RfxCas13d using a fluorescence reporter assay. **a** Schematic representation of the experimental design comparing glycyl tRNA–crRNA and ribozyme–crRNA arrays, each tested with GFP crRNA positioned at site 1 (GFP crRNA_P1), site 2 (GFP crRNA_P2), or site 3 (GFP crRNA_P3). **b** Green fluorescence micrographs of live RfxCas13d-expressing chicken cells imaged at 36 h post-transfection with crRNA array vectors. **c, d** Quantification of GFP-positive cells (c) and green mean fluorescence intensity (d) by flow cytometry. Data were analysed by one-way ANOVA relative to NT crRNA and GFP crRNA controls. Statistical significance is indicated by asterisks P < 0.0001 (****).

Fluorescence microscopy of DF1 cells transfected with the multiplex constructs revealed robust GFP knockdown across both expression strategies (Fig. 5b). However, positional analysis of GFP_crRNA activity indicated that ribozyme-based array achieved the most consistent GFP knockdown. When GFP_crRNA was positioned at site 1 or 2 within the ribozyme array, target suppression was maintained with only a modest (∼5%) reduction in activity compared to the single-guide construct (Fig. 5c, d). At position 3, GFP_crRNA retained full functionality in both array formats. In contrast, the glycyl tRNA-based array showed reduced target-specific activity when GFP_crRNA was expressed at positions 1 or 2.

These findings demonstrate that ribozyme-mediated crRNA processing enable more consistent guide activity across array positions, highlighting its suitability for reliable multiplexed crRNA delivery in RfxCas13d-based antiviral applications.

### 5. Multiplexed crRNA expression enhances antiviral potency against HPAIV in chicken cells

To determine whether crRNA multiplexing improves Rfx-Cas13d-mediated antiviral efficacy, RfxCas13d cell lines expressing PB1_M4 and NP_M10 individually or in combination (PB1_NP_M4+M10) were compared in two approaches using A/Chicken/Vietnam/08/2004[H5N1] (Fig. 6a). In the first approach, viral titres in infected RfxCas13d cell lines were quantified using TCID_50_ assays from supernatants collected at 24 and 48 h.p.i. At 24 h.p.i, cells expressing the PB1_NP_M4+M10 array exhibited significantly reduced viral titres (2.1 log) compared to single-guide cell lines (1.4 log reduction in NP_M10 and 1.3 log reduction in PB1_M4 cells), indicating enhanced antiviral activity through multiplexing (Fig. 6b). This enhanced antiviral effect persisted at 48 h.p.i, with both crRNA_M4 and crRNA_M4+M10 cell lines maintaining significantly lower titres relative to non-targeting controls (Fig. 6c). Corresponding cytopathic effects observed in the engineered DF1 cells are shown in Fig. S8.

**Fig. 6.**
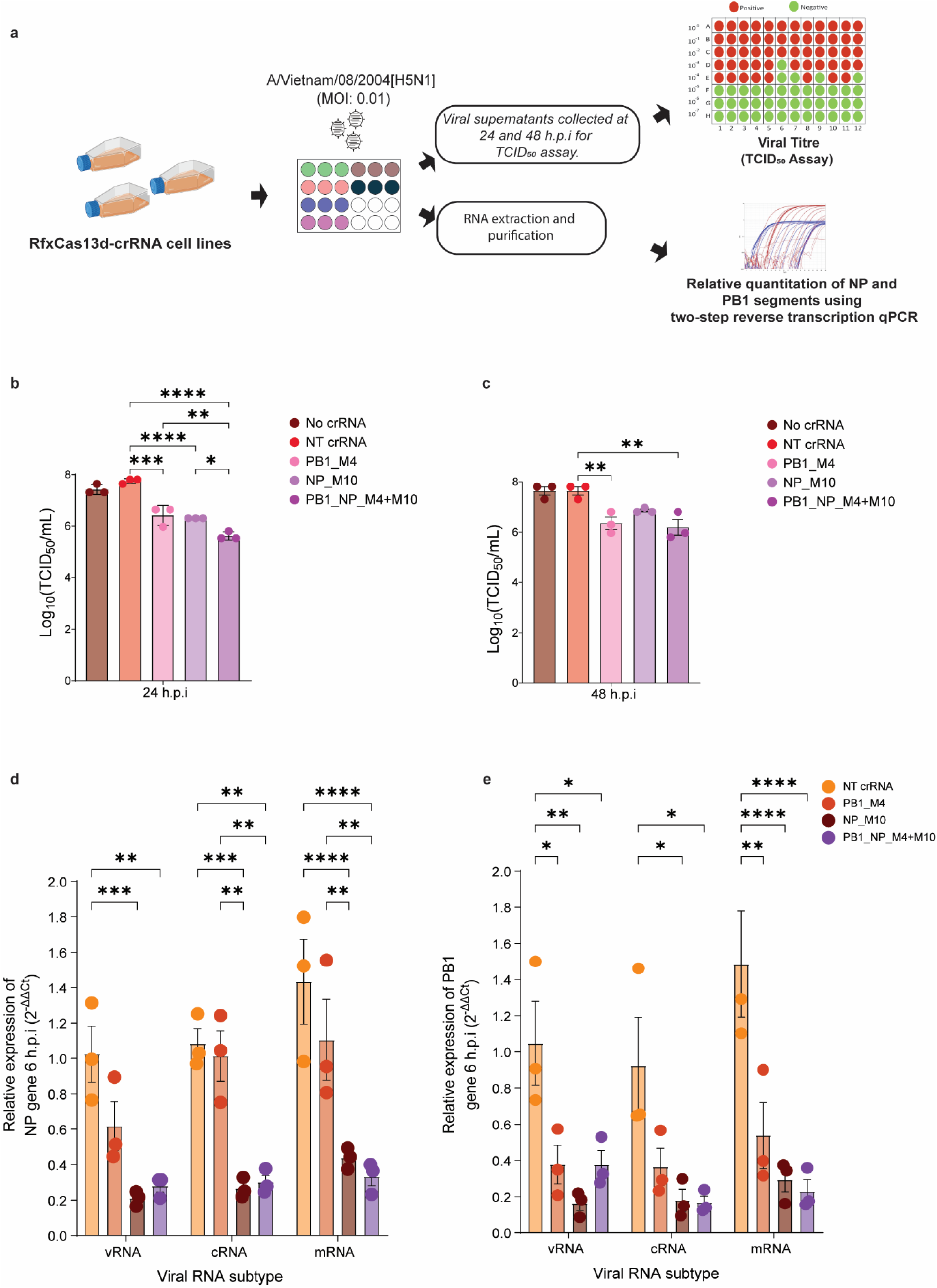
Comparative antiviral efficacy of single and multiplexed crRNA constructs targeting HPAIV. **a** Schematic overview of the experimental design used to evaluate the antiviral activity of RfxCas13d cell lines expressing either individual crRNAs (PB1_M4 or NP_M10) or a multiplexed PB1_NP_M4+M10 array against A/Chicken/Vietnam/08/2004[H5N1]. **b, c** Viral titres measured by TCID₅₀ assay in MDCK cells from supernatants collected at 24 h.p.i (b) and 48 h.p.i (c) from infected RfxCas13d cell lines expressing no crRNA, non-targeting (NT) crRNA, PB1_M4, NP_M10, or PB1_NP_M4+M10. **d, e** Relative quantification of viral RNA subtypes (vRNA, cRNA, and mRNA) for NP (d) and PB1 (e) segments at 6 h.p.i, determined by multiplexed TaqMan-based quantitative RT-qPCR from total RNA extracted from infected cells. Genomic vRNA levels were used as internal reference for normalization across RNA subtypes. Viral titres were calculated using the Reed–Muench method. Statistical analysis was performed using one-way ANOVA with NT crRNA as the reference group. Significance is indicated as follows: *p < 0.05; **p < 0.01; ***p < 0.001; ****p < 0.0001.

In the second approach, two step quantitative RT–PCR was performed on cellular RNA harvested from infected cells at 6 h.p.i to assess guide-specific knockdown of the target viral RNA intermediates. Results obtained from the relative qRT-PCR revealed that NP_M10 and the PB1_NP_M4+M10 array significantly reduced all viral RNA intermediates—vRNA, cRNA, and mRNA—of both NP and PB1 segments (Fig. 6d, 6e). In contrast, PB1_M4 selectively suppressed PB1-derived viral RNA intermediates, with negligible effect on NP RNA levels.

These findings emphasize the increased potential of crRNA multiplexing in enhancing RfxCas13d-mediated antiviral responses against HPAIV.

### 6. Broad-spectrum antiviral activity of RfxCas13d-crRNA arrays against circulating HPAIV H5N1 strains

Given the rapid evolution and global dissemination of highly pathogenic avian influenza viruses (HPAIVs), the development of broad-spectrum antiviral strategies is urgently needed. To assess the breadth of strain coverage and understand how broadly the tested guides can be used against circulating and emerging variants of H5N1 isolates, crRNA guide sequences PB1_M4 and NP_M10 were subjected to multiple sequence alignment across 16,919 HPAIV H5N1 genomes retrieved from the GISAID database. The bioinformatic pipeline for the analysis is shown in Fig. S9. From the study, the NP_M10 crRNA revealed to exhibit perfect sequence complementarity to 95.97% of isolates, while PB1_M4 matched 59.84% without mismatches. Interestingly, the combined coverage of both crRNAs reached up to 99.15% with at least no mismatch in one of the crRNAs, underscoring the potential of this dual-targeting strategy for broad-spectrum suppression of circulating H5N1 strains (Fig. 7a, b).

**Figure 7.**
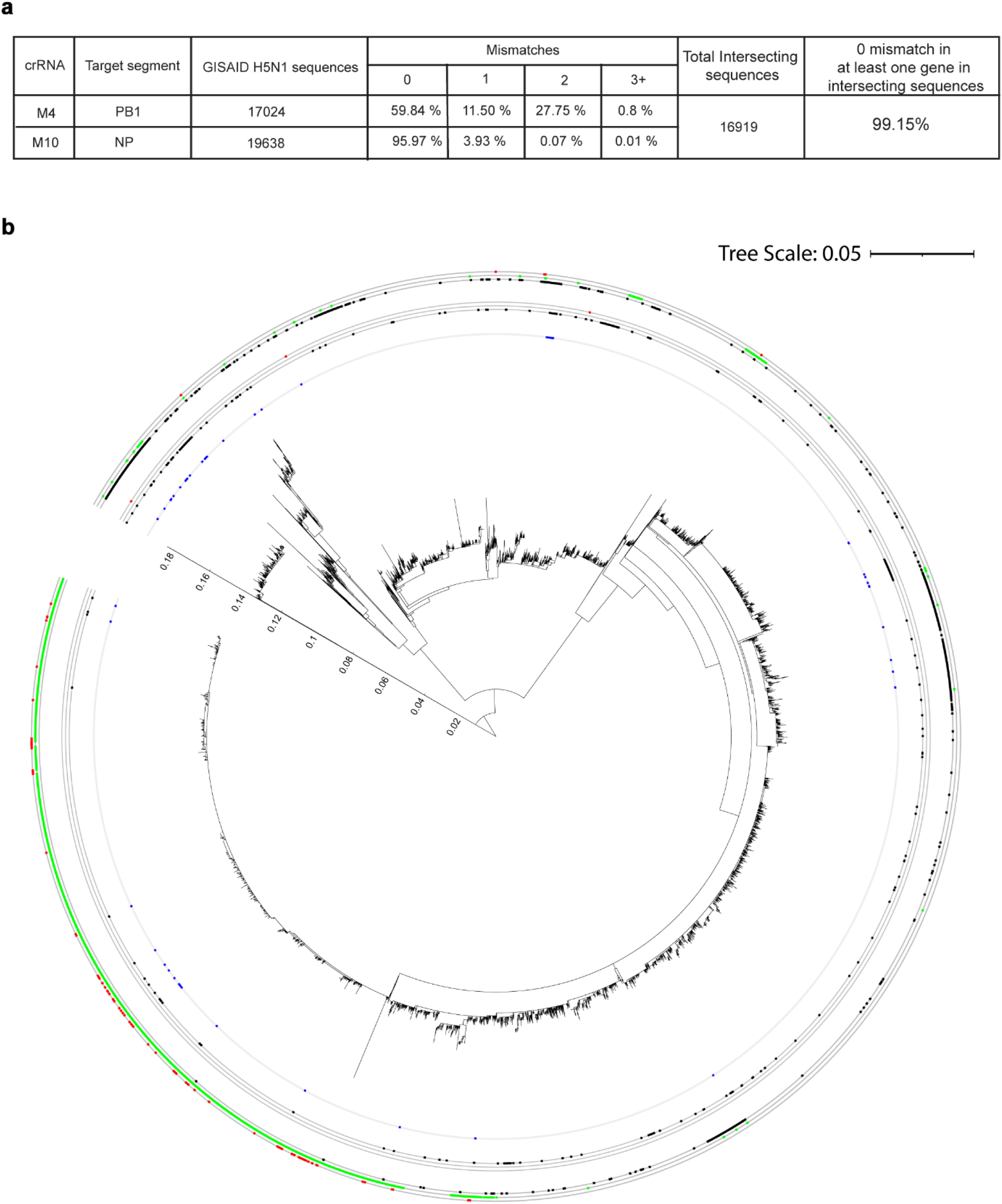
Conservation of crRNA target sites across highly pathogenic avian influenza H5N1 isolates. **a** Summary table showing mismatch frequencies at PB1_M4 and NP_M10 crRNA target sites across H5N1 highly pathogenic avian influenza virus (HPAIV) sequences retrieved from the GISAID database^33^. **b** Maximum-likelihood phylogenetic tree of H5N1 HPAIV isolates based on NP segment sequences retrieved from GISAID database, annotated with crRNA target site mismatches on PB1 and NP. Blue dots in the innermost annotation circle denote strains carrying at least one mismatch in both NP_M10 and PB1_M4 target sites; dots in the outer annotation circle indicate strains with 1 (inner ring, black dots), 2 (middle ring, green dots), or ≥3 (outer ring, red dots) mismatches within PB1_M4 target region; and dots in the middle annotation circle shows strains with 1 (inner ring, black dots), 2 (middle ring, green dots), or ≥3 (outer ring, red dots) mismatches within NP_M10 target region.

To evaluate the efficacy of RfxCas13d-mediated targeting against contemporary HPAIV strains, stable DF1 cell lines expressing RfxCas13d and individual or multiplexed crRNAs were challenged with A/Turkey/Indiana/22-003707-003/2022[H5N1], a clade 2.3.4.4b isolate. The schematic of methods employed to investigate the post -infection outcome is shown in Fig 8a. At 24 h.p.i, confocal immunofluorescence revealed diminished NP-associated green fluorescence in RfxCas13d cells expressing crRNAs PB1_M4 or NP_M10. Consistent with earlier observations, multiplexed crRNA (PB1_NP_M4+M10) produced the most pronounced suppression of NP levels (Fig. 8b). Consistent with the imaging data, viral titres at 24 h.p.i were significantly reduced in cells expressing single crRNAs targeting PB1 (PB1_M4) and NP (NP_M10), with reductions of 3.2 and 3.3 log units, respectively (Fig. 8c). Notably, the cell lines expressing the multiplexed crRNA array (PB1_NP_M4+M10) exhibited enhanced antiviral activity, achieving up to 4.39 log unit reduction in viral titre (Fig. 8c). By 48 h.p.i, PB1_M4 and the multiplex array-maintained suppression of viral replication, each showing approximately 1 log unit viral titre reduction, whereas NP_M10 alone did not confer measurable protection relative to non-targeting controls (Fig. 8d). Corresponding cytopathic effects shown in Fig. S10.

**Fig 8.**
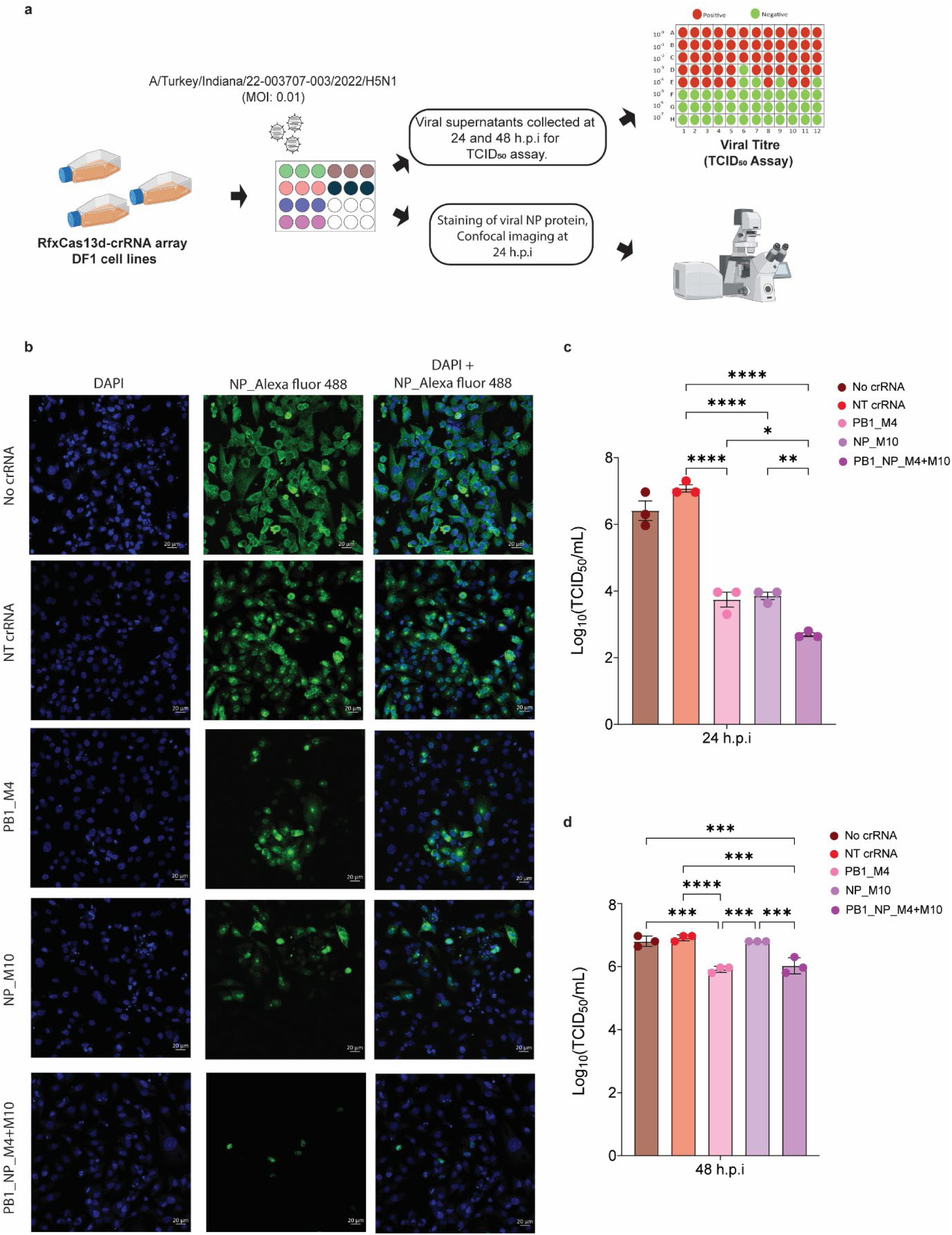
RfxCas13d-mediated antiviral activity against circulating H5N1 using positive sense RNA targeting crRNAs and the crRNA array. **a** Schematic of the workflow used to assess RfxCas13d-mediated inhibition of a currently circulating HPAI clade 2.3.4.4b virus. **b** Confocal immunofluorescence of RfxCas13d–crRNA-expressing cells infected with A/Turkey/Indiana/22-003707-003/2022[H5N1] (clade 2.3.4.4b) at 24 h post-infection, stained with anti-NP antibody (green) and counterstained with DAPI (blue). **c, d** Viral titres in supernatants from RfxCas13d-expressing cells at 24 h (**c**) and 48 h (**d**) post-infection, comparing No crRNA, NT crRNA, crRNA_M4, crRNA_M10, and crRNA_M4+M10 conditions. Titres were determined by the Reed–Muench method. Data are mean ± SEM; statistical significance was determined by one-way ANOVA: *p < 0.05, **p < 0.01, ***p < 0.001, ****p < 0.0001.

## Discussion

Programmable RNA-guided RNA-targeting systems have emerged as promising antiviral platforms against rapidly evolving pathogens such as influenza A viruses ^30–32^. Despite the potential, effective targeting of HPAIV strains remains challenging due to the extensive genetic plasticity of their RNA genomes ^14^. RNA interference (RNAi) strategies, including small interfering RNAs (siRNAs) or short hairpin (shRNA), have demonstrated potent antiviral activity against AIVs ^34,35^. However, RNAi-based antiviral strategies face several barriers, including the risk of the emergence of viral escape mutants, the requirement for high dosing to achieve robust inhibition, off-target gene silencing, dependence on host RNAi components, and the difficulty in achieving robust virus inhibition across different cellular compartments ^36^. In transgenic models, sustained expression of siRNAs or short hairpin RNAs (shRNAs) can overwhelm the RNAi machinery, disrupt endogenous microRNA networks, and in some cases induce host toxicity ^37^. Collectively, these constraints reinforce the need for alternative or complementary approaches to effectively restrict influenza replication. Recent studies have explored CRISPR/Cas13 systems for influenza inhibition in mammalian models ^30,32^, yet their performance varies widely across cell types and viral targets, complicating predictions of therapeutic utility. In avian transgenic models, to our knowledge only one *in vitro* study has assessed Cas13-mediated inhibition of influenza, showing limited efficacy against a lab-adapted, non-avian strain ^38^. Moreover, concerns over Cas13-induced collateral RNA degradation have raised questions about its safety in human applications, with some variants exhibiting cytotoxicity due to excessive off-target activity ^39,40^. Nonetheless, Cas13 enzymes with low to moderate collateral activity with or without host toxicity may hold promise for controlling HPAI outbreaks in poultry.

This study systematically evaluated five CRISPR/Cas13 systems for their target-specific and collateral RNA cleavage activities in chicken cells. Selected based on prior RNA-targeting studies in mammalian systems, five Cas13 variants were screened using a single crRNA targeting GFP, and a nuclear-localized expression strategy incorporating the Simian Virus 40 nuclear localization sequence ^28^. In our study, a nuclear localization signal was incorporated into the Cas13 constructs with a long-term goal to reflect upon the nuclear replication of influenza virus, thereby facilitating targeting of both positive- and negative-sense viral RNA species. While cytoplasmic mRNA targeting is limited, nascent viral mRNAs within the nucleus remain susceptible.

Among the tested Cas13 systems, RfxCas13d demonstrated superior target knockdown efficiency and robust expression, as indicated by red fluorescent protein reporter levels, suggesting high tolerability and functional activity in the chicken cells. Collateral activity profiling using a blue, fluorescent reporter revealed that RfxCas13d also exhibited the highest non-target RNA cleavage among the variants tested. Despite this elevated collateral activity, cell viability remained unaffected, consistent with previous findings^41^. These results highlight the potential of RfxCas13d as a well-tolerated and potent RNA-targeting tool in chicken cells. The lack of GFP knockdown by other Cas13 variants likely reflects the testing of a single GFP-targeting crRNA in single cell type rather than an intrinsic inactivity. As crRNA efficacy and spacer requirements vary across Cas13 variants, these results do not indicate non-functionality of other Cas13 variants but underscore the importance of crRNA design for each Cas13. Additionally, other factors such as insufficient protein expression of Cas13 may have contributed to this observed limited efficacy. Future studies assessing crRNA design and Cas13 expression levels will be essential to clarify these discrepancies. Given the strong knockdown activity of RfxCas13d, subsequent efforts focused on its application as an antiviral agent against influenza viruses.

A key advantage of CRISPR-based antiviral platforms is their capacity for multiplexing ^42^ which enables simultaneous targeting of multiple viral RNA regions using crRNA arrays. Although multiplexing has been extensively optimized for DNA targeting Cas9 systems, equivalent strategies for RNA targeting Cas13 platforms remain underexplored. To address this gap, we compared two established crRNA multiplexing methods previously applied in Cas9 systems-endogenous chicken glycyl tRNA ^43^ and self-processing ribozyme ^44^ sequences derived from hammerhead and hepatitis delta virus motifs. While both designs facilitated efficient reporter knockdown in RfxCas13d cells, ribozyme-based arrays consistently demonstrated superior performance, exhibiting robust activity across all crRNA positions within the multiplex. This enhanced efficacy likely reflects the precise and autonomous cleavage afforded by ribozyme motifs, which function independently of host RNase processing and are compatible with tissue specific promoters, thereby supporting flexible expression of crRNAs across cell types ^44^. To our knowledge, this study represents the first systematic comparison of crRNA multiplexing strategies for Cas13 systems. These findings establish ribozyme-based self-processing as a more efficient and versatile approach for RfxCas13d-mediated antiviral applications, providing a foundation for scalable RNA-targeting designs capable of addressing the genetic diversity of RNA viruses.

Influenza A virus, a member of the Orthomyxoviridae family, possesses a segmented genome comprising eight single-stranded, negative-sense RNA segments ^12^. As a Baltimore Class V virus, it replicates by synthesizing viral RNA intermediates from its negative-sense templates, with genome replication occurring entirely within the host cell nucleus. Maturation of pre-mRNA to mRNA enables translation of viral proteins, including those required for replication, ultimately leading to the formation of positive-sense copy ribonucleoproteins (cRNPs) that serve as templates for generating progeny negative-sense vRNPs. The viral polymerase complex, composed of PB2, PB1, and PA alongside the nucleoprotein (NP), plays a central role in replication and transcription ^45^.

Our analyses reveal that the antiviral efficacy of RfxCas13d is strongly influenced by crRNA strand orientation and viral replication dynamics. Positive-sense targeting crRNAs exert broader and more potent antiviral effects by concurrently reducing multiple viral RNA intermediates, whereas negative-sense targeting preferentially impacts genomic RNA with more limited downstream effects. These strand-dependent differences are further shaped by segment-specific replication and transcription kinetics, highlighting the importance of RNA accessibility in determining RfxCas13d activity. Together, our findings define key parameters governing RfxCas13d-mediated antiviral suppression and provide a mechanistic framework for the rational design of crRNAs to maximise antiviral efficacy against highly pathogenic avian influenza viruses.

In this study, crRNAs directed against PB1, and NP’s positive-sense RNA significantly suppressed viral replication and conferred prolonged protection in chicken cells, outperforming those targeting negative-sense RNA. Mechanistically, the positive sense RNA targeting guides inhibited all the viral RNA species including the negative sense RNA strand in our study. The enhanced activity is likely driven by the greater abundance of positive-sense RNA species. Our long-read sequencing analysis showed that, across all samples, reads corresponding to cRNA were approximately twofold higher than those mapping to viral genomic RNA. In addition, the presence of nascent viral mRNAs in the nucleus provides multiple accessible targets when directing Cas13d against positive-sense RNA, potentially amplifying collateral activity. This increased targeting landscape, comprising both pre-mRNA and cRNA in the nucleus, is consistent with recent findings that higher RfxCas13d expression is associated with increased collateral activity ^46^. Because cRNAs play an early and active role in replication and directly contribute to generating negative-sense RNA, their disruption may explain the observed reduction in vRNA levels. Collateral impact on other viral RNA segments and intermediates further contributed to lowering viral titres. Moreover, targeting pre-mRNA also disrupts viral protein synthesis over an extended period, reinforcing the antiviral effect also reducing the replication of negative sense RNA. To our knowledge, this is the first study to directly compare crRNAs targeting positive-sense RNA versus negative-sense RNA intermediates in cells infected with highly pathogenic avian influenza viruses. Nonetheless, prior work in other influenza strains (excluding HPAIVs) has demonstrated that Cas13-mediated targeting can effectively reduce viral RNA levels, reiterating the broader potential of this approach ^30,31^. A key distinction between previous studies^30,31^ and ours is the use of transgenic cell lines engineered to express antiviral RfxCas13d-crRNAs, with the goal of evaluating their potential in transgenic animals, while previous studies were more focused in delivering the synthetic RNAs in human cells. Additional differences include the initial viral inoculum (lower MOI^31^), the targeted viral RNA segments (PB1^30^ and PB2^30,31^), the specific Cas13 variants used (LbuCas13a) ^30,31^, and the incorporation of chemical modifications to enhance guide RNA stability ^31^. Furthermore, as the viral subtypes examined in those studies differ from the highly pathogenic avian influenza viruses (HPAIVs) used here, direct comparison of outcomes may not be appropriate. Notably, similar trends were observed with respect to targeting positive-versus negative-sense viral RNA. Additionally, the current design localizes RfxCas13d to the nucleus, which focuses its activity against viral RNAs in the nucleus directly targeting the viral RNA intermediates. Future improvements could involve developing Cas13 constructs capable of targeting viral RNA in both the cytoplasm and nucleus, thereby enhancing antiviral efficacy.

Given that birds serve as natural reservoirs for highly pathogenic avian influenza (HPAI) viruses ^12^, we evaluated the antiviral efficacy of RfxCas13d in chicken cells. Three HPAI strains were selected for testing: two from the H5 lineage and one from the H7 lineage. The H5N1 strain A/Chicken/Vietnam/08/2004 ^47^ was chosen for its rapid replication kinetics, zoonotic potential, and documented 100% mortality in avian hosts. In contrast, the H7N7 strain A/Chicken/Lethbridge/09/2020 ^48^ represents a slower-replicating virus yet remains lethal to chickens and was implicated in a major HPAI outbreak in Australia. RfxCas13d-expressing cells equipped with crRNAs targeting positive-sense viral RNA demonstrated significant inhibition of both HPAI strains. Notably, viral titres of H5N1 were reduced by up to 1.4 log units in the most effective crRNA line, whereas H7N7 replication was suppressed below the limit of detection for 48 h.p.i. These findings suggest that viral replication kinetics may critically influence the antiviral potency of RfxCas13d.

Our findings also demonstrate that enhancing RfxCas13d antiviral performance through crRNA multiplexing represents a promising strategy for combating genetically diverse HPAI viruses. By incorporating self-cleaving ribozyme sequences, we achieved coordinated expression of multiple crRNAs within a single construct, enabling simultaneous targeting of distinct viral RNA regions. This multiplexing approach consistently produced greater viral inhibition compared to single crRNA treatments, particularly when directed against positive-sense viral RNA, aligning with our earlier observations that such targets yield superior Cas13-mediated degradation. Importantly, when tested against two H5-lineage viruses, including the currently circulating clade 2.3.4.4b strain A/Turkey/Indiana/22-003707-003/2022[H5N1], multiplexed crRNAs markedly enhanced viral suppression. The observed increase in antiviral activity using the multiplexed array highlights the enhanced effect of targeting multiple viral RNA intermediates simultaneously.

Beyond improved potency, multiplexing may also offer a critical advantage in mitigating viral escape. Retrospective sequence analyses revealed that the selected crRNAs target highly conserved regions across diverse H5N1 genomes, suggesting that this design could confer broad-spectrum activity against circulating and emerging variants. Together, these results demonstrate the potential of multiplexed RfxCas13d systems as a next-generation antiviral platform capable of both amplifying efficacy and maintaining resilience against the genetic plasticity of rapidly evolving avian influenza viruses.

A critical challenge for the sustainability of these advancements lies in their effective delivery across both human and animal systems. In this study, we focused on generating transgenic cell lines with the long-term goal of developing disease-resilient poultry. Transgenic animals could offer a promising solution to limiting viral replication within hosts, thereby reducing the risk of zoonotic spillover. However, societal acceptance of genetically modified animals remains a significant barrier. Emerging technologies, particularly synthetic RNA platforms, provide renewed hope for future generations. Nevertheless, combating highly pathogenic viruses such as HPAIV will require stronger regulatory support and endorsement from government agencies to translate these innovations into practical solutions.

To further contextualize the significance of this work, it is important to consider its alignment with the interconnectedness of animal, human, and environmental health. Highly pathogenic avian influenza (HPAI) viruses, particularly those of the H5 lineages, continue to pose a substantial threat due to their zoonotic potential and rapid genomic evolution. By demonstrating the efficacy of RfxCas13d in suppressing HPAI replication in chicken cells, this study offers a targeted approach to reducing viral burden in avian reservoirs as an essential step in mitigating the risk of cross-species transmission. The development of scalable, precision-guided antiviral strategies for poultry not only strengthens biosecurity within agricultural systems but also contributes to global pandemic prevention efforts. Incorporating CRISPR-based tools into surveillance and outbreak response infrastructures may represent a pivotal advancement in proactive disease control, reinforcing the critical role of veterinary interventions in safeguarding public health.

In summary, this study establishes RfxCas13d as a highly effective and well-tolerated RNA targeting antiviral platform in chicken cells, capable of suppressing multiple HPAI strains with minimal cytotoxicity. Future work integrating *in vivo* validation, expanded strain coverage, evaluation of emergence of escape mutants and optimized delivery systems will be critical to translating this platform into field-ready antiviral interventions.

## Methods

### 1. Construction of Cas13 and crRNA Expression Vectors for Functional Screening

To evaluate target-specific and collateral activity across multiple Cas13 systems, unbiased expression constructs were generated using a Tol2 transposon backbone. The Cas13 expression vector comprised the following elements: Tol2LHE–ICeuI–Cbh promoter–NotI–nuclear localisation signal (NLS)–Cas13 (LwaCas13a, PspCas13b, RfxCas13d, Cas13e3 or Cas13x(HF))–NLS–PacI–IRES–miRFP670nano–SV40 polyA–Tol2RHE (for vector map refer to Fig. S1). Protein sequences of Cas13 variants used in this study are provided in Table S1. A corresponding crRNA expression vector was designed with ICeuI–human U6 promoter–BbsI–BbsI–CMV promoter–mTagBFP2–SV40 polyA (for vector map refer to Fig. S2). Nucleotide sequences of both the backbone vectors for Cas13 and crRNA are provided in Table S2. Fluorescent markers were used to facilitate selection: miRFP670nano (red) for Cas13 constructs and mTagBFP2 (blue) for crRNA vectors. Synthetic crRNA gene blocks were designed and cloned into the BbsI sites of the crRNA vector. Direct repeat (DR) sequences specific to each Cas13 variant, along with spacer sequences used in this study, are listed in Table S3.

For genomic integration, the crRNA cassette comprising the human U6 promoter, crRNA, and mTagBFP2 marker was subcloned into the Cas13 Tol2 backbone between the *ICeu*I and *Fse*I restriction sites. Additionally, a GFP expression vector (CAAGS-GFP-polyA) was employed as a target construct for initial screening of Cas13-mediated target knockdown and collateral activity (Fig. S1 and Fig. S2).

### 2. Cell culture

Chicken embryonic fibroblast cells (DF1) used throughout the study were maintained in modified Dulbecco’s Modified Eagle Medium (DMEM) supplemented with 2 mM L-glutamine, 4,500 mg/L glucose, 1 mM sodium pyruvate, 1,500 mg/L sodium bicarbonate, 10 mM HEPES buffer, 100 U/mL penicillin, 100 µg/mL streptomycin, and 2.5 µg/mL amphotericin B, with or without 10% foetal bovine serum (FBS), depending on experimental requirements. Madin-Darby Canine Kidney (MDCK) cells were cultured in Eagle’s Minimum Essential Medium (EMEM) supplemented with Earle’s Balanced Salt Solution, non-essential amino acids, 2 mM L-glutamine, 1 mM sodium pyruvate, 1,500 mg/L sodium bicarbonate, 100 U/mL penicillin, and 100 µg/mL streptomycin. When required, 10% FBS was added to the culture medium. All cell lines were incubated at 37 °C in a humidified atmosphere containing 5% CO₂.

### 3. Comparative Functional Analysis of Cas13 Variants in DF1 Cells

To compare the target-specific activity of five Cas13 variants—LwaCas13a, PspCas13b, RfxCas13d, Cas13e3 and Cas13x(HF)—variant-specific direct repeat (DR) sequences were incorporated into crRNA constructs alongside a previously validated GFP-targeting spacer sequence. The positioning of DR elements within each crRNA was determined according to the structural preferences reported for each Cas13 system in prior studies ^28,41,49,50^. For each Cas13 variant, a corresponding non-targeting crRNA (NT crRNA) was also designed and cloned into the crRNA expression vector to serve as a negative control.

#### 3.1 Transfection of DF1 Cells with Cas13, crRNA, and GFP Reporter Constructs

Wild-type DF1 cells were seeded in 24-well plates at 2.0 × 10⁵ cells/well and incubated overnight at 37 °C with 5% CO₂. After washing with PBS, cells were prepared for transfection using 300 µL Opti-MEM (Thermo Fisher Scientific) supplemented with insulin, transferrin, hypoxanthine, thymidine, trace elements, and 2.4 g/L sodium bicarbonate. Plasmid DNA (Cas13, crRNA, GFP vectors in a 9:9:2 ratio; 1 µg total) was diluted Opti-MEM. Lipofectamine 2000 was prepared per manufacturer’s instructions and incubated for 5 min at room temperature. Equal volumes of DNA and Lipofectamine were combined and incubated for 15–20 min to form complexes, which were added to cells and incubated at 37 °C with 5% CO₂. After 2–4 h incubation, cells were washed with PBS and cultured in DMEM with 10% FBS for 2–3 days.

#### 3.2 Microscopy

Expression of the Cas13 vector was validated by monitoring red fluorescent protein (miRFP670nano) in live DF1 cells using the Sartorius Incucyte® S3 live-cell imaging system at 36 h post-transfection. Target-specific activity of Cas13 variants was assessed by quantifying GFP knockdown in the same imaging platform. Collateral activity was evaluated by detecting blue fluorescence (mTagBFP2) from the crRNA vector using a Leica epifluorescence microscope.

#### 3.3 Flow cytometry

Following 36 h of incubation and live-cell imaging, DF1 cells were harvested using 0.05% Trypsin–EDTA solution, washed with phosphate-buffered saline (PBS), and resuspended in pre-warmed DMEM supplemented with 10% foetal bovine serum (FBS). Cells were transferred to 96-well U-bottom plates for flow cytometric analysis using the BD LSR Fortessa II cell analyser.

Single-cell populations were gated using forward and side scatter plots. Fluorescence signals were acquired as follows: red fluorescence (miRFP670nano) using a 633 nm red laser for excitation and a 660/20 bandpass filter for emission; green fluorescence (GFP) using a 488 nm blue laser, 505 LP dichroic mirror, and 530/30 bandpass filter; and blue fluorescence (mTagBFP2) using a 405 nm violet laser and a 450/50 bandpass filter. Compensation for spectral overlap was performed using single-colour control samples. As red fluorescence marked Cas13 vector expression, green and blue fluorescence signals were resolved specifically within the red-positive population. Flow cytometry data were analysed using the latest version of FlowLogic software. Gating strategy applied to analyse the raw data is shown in Fig. S11.

### 4. Engineering of DF1 cell lines to stably express RfxCas13d

To generate stable RfxCas13d-expressing chicken fibroblast cell lines, DF1 wild-type cells were co-transfected with the RfxCas13d expression vector with Tol2 backbones and a transposase vector at a 1:2 molar ratio. Transfection was performed as described in above in section 3.1 and cells were incubated 5–10 days at 37 °C in a humidified atmosphere containing 5% CO₂ to allow genomic integration. Fluorescence-assisted cell sorting (FACS)

Transfected DF1 cells were harvested using 0.05% trypsin–EDTA, washed with phosphate-buffered saline (PBS), and resuspended in complete DMEM. Expression of the red fluorescent marker miRFP670nano, encoded by the RfxCas13d vector, was used to identify transgene-positive cells and sorted using BD FACS Aria II. Cells expressing miRFP670nano (excitation: 645 nm; emission: 670 nm) were resolved using a 633 nm red laser and a 660/20 bandpass filter on BD FACSAria II (gating strategy provided in Fig. S12). A total of 2 × 10⁵ red fluorescent DF1 cells were sorted and seeded into 24-well plates for expansion. Cultures were subsequently transferred to 75 cm² tissue culture flasks and grown to confluency. To further enrich the transgenic population, a second round of sorting was performed, yielding 5 × 10⁵ purified RfxCas13d-expressing cells, which were seeded into 25 cm² flasks. The purified cell lines were maintained, stored and used further.

### 5. Design, Engineering, and Functional Evaluation of RfxCas13d-Based Antiviral Strategies Against Influenza A Virus

#### 5.1 Computational Design of Cas13 crRNAs Targeting Influenza A Virus

A computational pipeline was developed to facilitate Cas13 guide RNA (crRNA) design against influenza A virus. Three viral gene segments, nucleoprotein (NP), polymerase basic protein 1 (PB1), and polymerase basic protein 2 (PB2), were selected as representative targets based on their essential roles in viral replication. Viral genome sequences from H1N1, H5N1 and H7N7 subtypes were analysed to identify conserved 24-nucleotide regions across the influenza A genome. Conservation was assessed via multiple sequence alignments, enabling prioritisation of candidate regions with broad subtype coverage.

For each candidate guide, a comprehensive set of sequence-derived features was computed, including nucleotide composition, predicted RNA secondary structure, minimum free energy (MFE), homopolymer runs, GC content, mismatch tolerance, and genomic context (coding sequence or untranslated region). These features, along with local sequence context, were used to train a convolutional neural network (CNN) implemented in Python (v3.10) using TensorFlow (v2.12). The model was trained using a cross-entropy loss function and optimised with the Adam algorithm. Hyperparameters were tuned via five-fold cross-validation. Model performance was evaluated using Pearson correlation, precision, recall and F1 score metrics to ensure generalisability and robustness of guide efficacy predictions across viral subtypes.

#### 5.2 Screening of Positive-Sense RNA-Targeting crRNAs Against Influenza A Virus

To experimentally screen crRNAs targeting positive-sense viral RNA, nine crRNAs were designed against conserved regions of the PB2 (PB2_M1–M3), PB1 (PB1_M4–M6), and NP (NP_M7–M9) segments. Each crRNA was cloned into the BbsI restriction sites of the Tol2 backbone crRNA expression vector. Successful insertion of crRNAs was confirmed by Sanger sequencing. Nucleotide sequences of the crRNAs provided in Table S4, crRNA design strategy provided in Fig. S4.

DF1 cells stably expressing RfxCas13d were seeded in 24-well plates at a density of 2 × 10⁵ cells per well and cultured at 37°C with 5% CO₂. Following overnight incubation, cells were transfected in triplicate with 900 ng of crRNA vector per well and maintained under identical conditions. After 24 h, transfected cells were washed with phosphate-buffered saline (PBS) and supplemented with 250 µL of DMEM lacking foetal bovine serum. The A/WSN/1933[H1N1] virus was diluted to achieve a multiplicity of infection (MOI) of 0.01 and used to infect the transfected DF1 cells. Infected cultures were incubated for 24 h, washed twice with PBS, and total RNA was extracted using the RNeasy Mini Kit (Qiagen) according to the manufacturer’s protocol.

Quantification of viral replication was performed by measuring M-gene expression using the AgPath-ID™ One-Step RT-PCR reagents on the QuantStudio™ 3 Real-Time PCR System. Relative expression levels were calculated using the 2^−ΔΔCt^ method, normalising M-gene Ct values to endogenous 18S rRNA controls from both infected and uninfected samples. The Panflu Primers and probe sequences used in the assay are listed in Table S7.

#### 5.3 Functional Evaluation of RfxCas13d-Expressing DF1 Cell Lines Targeting Influenza A Virus

##### 5.3.1 Engineering Stable RfxCas13d DF1 Cell Lines Expressing Influenza Virus-Targeting crRNAs

To generate stable RfxCas13d-expressing cell lines targeting influenza virus RNA, DF1 cells were further engineered to co-express crRNAs specific to either positive- or negative-sense viral RNA. Based on initial screening results, three crRNAs targeting positive-sense RNA (PB1_M4, PB1_M6 and NP_M7) were selected and cloned into Tol2 backbone vectors. These constructs were co-transfected with a transposase-expressing vector into RfxCas13d cell lines to facilitate genomic integration.

In parallel, five crRNAs targeting negative-sense RNA (NP_G1–G3 and PB1_G4-G5) were designed, cloned into Tol2 vectors, and transfected into RfxCas13d cells to establish five additional crRNA-expressing lines. Of these, NP_G2, NP_G3 and PB1_G5 exhibited perfect sequence complementarity to the negative-sense RNA of the A/WSN/1933[H1N1] strain, while NP_G1 and PB1_G4 contained natural mismatches against A/WSN/1933[H1N1] virus but were perfect match for HPAI sequences (see Table S4 and Table S5). Notably, NP_G1 was reverse complementary to the NP_M7 in highly pathogenic influenza strains.

From the initial panel of A/WSN/033[H1N1] specific positive-sense RNA-targeting crRNAs, two guides demonstrating the highest efficacy were selected for further optimisation. These guides (PB1_M4 and NP_M7) were redesigned or reused to target conserved regions in three highly pathogenic avian influenza viruses: A/Chicken/Vietnam/08/2004[H5N1] (PB1_M4 and NP_M10), A/Chicken/Lethbridge/09/2020[H7N7] (NP_M10 and PB1_M11) and A/Turkey/Indiana/22-003707-003/2022[H5N1] (Clade 2.3.4.4b) (PB1_M4 and NP_M10). Redesigned crRNAs were cloned into Tol2 backbone vectors and transfected into RfxCas13d cell lines. Nucleotide sequences for the crRNA spacers are listed in Table S5.

DF1 cells co-expressing both red and blue, fluorescent markers were sorted using FACS Aria II and cultured, achieving a minimum purity threshold of 90% prior to downstream infection assays. Gating strategy applied for sorting the RfxCas13d crRNA cell lines provided in Fig. S13.

##### 5.3.2 Antiviral assessment of RfxCas13d-crRNA DF1 cell lines against influenza A virus

Purified RfxCas13d-crRNA cells were evaluated for antiviral activity against influenza A viruses. All infections involving highly pathogenic avian influenza strains were conducted within physical containment level 3 (PC3) facilities in accordance with institutional biosafety protocols. For each assay, three replicates of engineered DF1 cells were seeded at the standard density in 24-well tissue culture plates and infected at a multiplicity of infection (MOI) of 0.01 in DMEM lacking foetal bovine serum, irrespective of viral strain. Cultures were incubated for up to 72 h.p.i or until cytopathic effects (CPEs) were observed.

Viral supernatants were harvested every 24 h.p.i, with earlier collection if complete CPE observed prior to the 72-hour endpoint. Phase-contrast microscopy was performed at 24-hour intervals to monitor CPE, and representative images were captured for each time point.

##### 5.3.3 Quantification of Influenza Viral Load Using TCID₅₀ Assay

Viral titres in supernatants collected from infection assays were quantified using the tissue culture infectious dose 50 (TCID₅₀) assay in Madin-Darby Canine Kidney (MDCK) cells. MDCK cells were seeded into 96-well plates at a density of 2 × 10⁴ cells per well in Eagle’s Minimum Essential Medium (EMEM) supplemented with 10% foetal bovine serum (FBS) and incubated overnight at 37 °C with 5% CO₂. Following incubation, cells were washed with phosphate-buffered saline (PBS) and replenished with EMEM lacking FBS.

Virus supernatants were subjected to 10-fold serial dilutions up to 10⁻⁷, and each dilution was applied to MDCK cells in at least four technical replicates. After 72 h.p.i, cytopathic effects (CPEs) were assessed microscopically, and viral titres were calculated using the Reed–Muench method, expressed as Log₁₀TCID₅₀/ml for each sample.

##### 5.3.4 Direct cDNA Sequencing and Strand-Resolved Analysis of Influenza Viral RNA Species and Cas13 Cleavage Sites

Total RNA was extracted from H5N1-infected cells and separated into polyadenylated (mRNA) and non-polyadenylated RNA fractions using the Dynabeads™ mRNA Purification Kit (Thermo Fisher Scientific, cat. no. 61006) according to the manufacturer’s instructions. The mRNA fraction was used directly for library preparation using the Direct cDNA Sequencing Kit (Oxford Nanopore Technologies; SQK-DCS109) ^51^, barcoded with the Native Barcoding Expansion Kit (EXP-NBD104/114), and sequenced on MinION flow cells (R9.4.1) to generate long-read mRNA sequences.

The non-polyadenylated RNA fraction was purified using the RNeasy MinElute Cleanup Kit (Qiagen), followed by enzymatic 3′-end polyadenylation ^52^. Polyadenylated non-mRNA species were subsequently processed using the same direct cDNA sequencing library preparation workflow as for the mRNA fraction, using 4 ng cDNA input per barcoded sample. Sequencing libraries were loaded onto MinION flow cells and sequenced according to the manufacturer’s recommendations.

Raw signal data (Pod5) were base-called using Dorado (v1.3.0) ^53^ with the high-accuracy model (dna_r9.4.1_e8_hac@v3.3). Basecalling was performed using a minimum quality score (Q-score) cutoff of 7. The resulting reads were demultiplexed into sample-specific FASTQ files using the dorado demux tool, targeting the EXP-NBD104/114 barcoding kit. To preserve full-length sequence integrity, no-trimming mode was applied during the basecalling process.

Base-called Oxford Nanopore long-read direct cDNA sequences were aligned to to the A/H5N1 reference genome using minimap2 (v2.3) with the -ax map-ont preset. The resulting alignments were converted to BAM format, sorted, and indexed using SAMtools (v1.23) ^54^. Mapping statistics, including total mapped reads and per-segment coverage, were extracted using the samtools idxstats command. Alignment summaries were generated for each barcode to quantify the distribution of reads across the viral segments.

##### 5.3.4.1 Viral RNA species differentiation and quantification

To quantify distinct viral RNA species, aligned reads were categorized into mRNA, cRNA, and vRNA using a strand-aware computational pipeline. Mapping statistics were extracted using Rsamtools (v2.22.0) ^55^ within the R statistical environment (v4.4.0) ^56^. For mRNA-enriched libraries, total viral mapping was calculated across all eight genomic segments. For non-mRNA libraries, reads were further partitioned by strand orientation where sense-strand reads (positive strand) were designated as cRNA, while anti-sense reads (negative strand) were designated as vRNA. Segment-specific abundance was calculated as a proportion of total sequenced reads (mapped + unmapped) to normalize for library size across barcodes. Statistical significance between experimental groups and non-targeting controls was determined using pairwise t-test with Benjamini-Hochberg (BH) p-value adjustment for multiple comparisons. Quantitative visualizations, including segment-wise distribution plots and overall viral proportion bars, were generated using ggplot2 ^57^.

##### 5.3.4.2 Cas13 cleavage site analysis

Reads mapping to the viral nucleoprotein (NP) segment were extracted for downstream analyses. Putative Cas13-mediated cleavage sites were inferred from soft-clipped read termini encoded in the CIGAR strings of aligned reads ^58^. Cleavage coordinates were calculated relative to the reference genome while accounting for read orientation. Cleavage positions from individual reads were aggregated to identify recurrent cleavage events along the NP segment.

#### 5.3.5 Transcript-Level Evaluation of crRNA Activity Against Influenza Viral RNA Subtypes

To investigate the differential activity of crRNAs targeting positive- and negative-sense viral RNA, crRNAs were designed to target distinct regions within the PB1 segment of the A/Chicken/Vietnam/08/2004[H5N1] strain. RfxCas13d-expressing DF1 cell lines expressing either a non-targeting control crRNA, crRNA_M4 (positive-sense target), or crRNA_G5 (negative-sense target) were seeded into 6-well plates at a density of 1 × 10⁶ cells per well in complete DMEM containing 10% foetal bovine serum (FBS), with three biological replicates per condition. After overnight incubation at 37 °C and 5% CO₂, cells were washed with phosphate-buffered saline (PBS), switched to serum-free DMEM, and infected with A/Chicken/Vietnam/08/2004[H5N1] at a multiplicity of infection (MOI) of 0.01. The infection medium was incubated with cells for 6 h, followed by PBS washes and total RNA extraction using the RNeasy Plus Mini Kit (Qiagen), according to the manufacturer’s protocol.

To quantify viral RNA subtypes, genomic RNA (vRNA), complimentary RNA (cRNA), and messenger RNA (mRNA)—a strand specific two-step reverse transcription quantitative PCR (RT-qPCR) assay was performed ^59^. First-strand cDNA synthesis was carried out using Superscript IV reverse transcriptase (Thermo Fisher Scientific Aust Pty Ltd) with gene-specific primers for vRNA and cRNA in separate reactions. mRNA-derived cDNA was generated using anchored oligo(dT) primers (Integrated DNA Technologies). An 18S rRNA-specific reverse transcription primer was included as an endogenous control for all reactions (Thermo Fisher Scientific Aust Pty Ltd). Quantitative PCR was performed using target-specific TaqMan probes and primer sets (Table S6 and Table S7). A Cy5-labelled probe was used for PB1_M4 target region, a FAM-labelled probe for PB1_G5, and a VIC-labelled probe for 18S rRNA quantification. Reactions were set using TaqMan Fast Advanced Master Mix (Thermo Fisher Scientific Aust Pty Ltd) on the QuantStudio™ 6 Pro Real-Time PCR System, and relative PB1 expression levels were analysed using the 2^−ΔΔCt^ method.

### 6. Multiplexed crRNA array engineering and functional validation in RfxCas13d-expressing chicken cells

#### 6.1 Design and functional screening of multiplexed crRNA arrays

Multiplexed crRNA arrays for RfxCas13d were engineered using two strategies adapted from Cas9 systems. In the first, chicken glycine tRNA (glytRNA) sequences sourced from RNA central were inserted between crRNAs to promote endogenous processing. In the second, hammerhead (HH) and hepatitis delta virus (HDV) ribozymes were positioned at the 5′ and 3′ ends of each crRNA, respectively, to enable autonomous cleavage. Nucleotide sequences of chicken glycyl tRNA and the ribozymes are listed in Table S8.

To assess positional effects on crRNA activity, GFP-targeting crRNA (GFP crRNA) and a non-targeting control were assembled into arrays containing crRNA1 at position 1, 2, or 3, flanked by non-specific crRNAs. Constructs were designated glytRNA_GFP crRNA_Pn or ribozyme_GFP crRNA_Pn, where n indicates GFP crRNA position. Nucleotide sequences of multiplexed crRNAs using in this study are listed in Table S9.

Synthetic gene blocks (Invitrogen GeneArt, Thermo Fisher Scientific) were cloned into the BbsI site of the crRNA expression vector and verified by Sanger sequencing. RfxCas13d DF1 cells were co-transfected with GFP and crRNA array plasmids at a 1:9 ratio and incubated for 36 h. GFP knockdown was quantified by live-cell imaging (Incucyte® S3, Sartorius) and flow cytometry (Fortessa LSR II, BD), with data analysed using FlowLogic (latest version).

#### 6.2 Comparative Evaluation of Single and Multiplexed crRNAs Targeting HPAIVs

To assess the antiviral efficacy of multiplexed crRNA arrays relative to individual crRNAs, DF1 cell lines stably expressing RfxCas13d and either PB1_M4, NP_M10, PB1_NP_M4+M10, a non-targeting control (NT crRNA), or no crRNA were seeded into two 24-well plates and one 6-well plate. Cells were incubated overnight at 37 °C in a humidified atmosphere containing 5% CO₂. Nucleotide sequence of the multiplexed crRNA PB1_NP_M4+M10 is listed in Table S9.

In the 24-well format, cells were infected with highly pathogenic avian influenza viruses (HPAIVs), A/Chicken/Vietnam/08/2004[H5N1] and A/Turkey/Indiana/22-003707-003/2022[H5N1], at a multiplicity of infection (MOI) of 0.01. Supernatants were collected at 24 and 48 h.p.i for viral titre quantification using the TCID₅₀ assay. Cytopathic effects (CPEs) were monitored microscopically, and viral titres were calculated using the Reed–Muench method.

In parallel, cells seeded in 6-well plates were infected with HPAI virus A/Chicken/Vietnam/08/2004[H5N1] and incubated for 6 hours. Following infection, cells were washed with PBS, and total RNA was extracted using the RNeasy Plus Mini Kit (Qiagen), according to the manufacturer’s protocol. To evaluate crRNA-mediated effects on viral RNA subtypes, a two-step reverse transcription quantitative PCR (RT–qPCR) assay was performed as described in Section 5.3.4.

Multiplexed qPCR assays were performed using TaqMan Fast Advanced Master Mix (Thermo Fisher Scientific Aust Pty Ltd) on the QuantStudio™ 6 Pro Real-Time PCR System. Fluorophore-labelled probes were used to detect PB1 (Cy5), NP (FAM), and 18S rRNA (VIC), alongside corresponding forward and reverse primers flanking each probe-binding site. Raw fluorescence data were processed to quantify relative gene expression across conditions.

#### 6.3 Confocal Immunofluorescence Detection of Viral nucleoprotein (NP)

Chicken cells were seeded on coverslips in 24-well plates and infected with A/Turkey/Indiana/22-003707-003/2022[H5N1] at a multiplicity of infection (MOI) of 0.01. At 24 h post-infection, cells were fixed with 4% PFA, permeabilized with 0.1% Triton X-100, and blocked with 3% bovine serum albumin. Cells were incubated overnight at 4°C with mouse anti-NP antibody, followed by Alexa fluor 488-conjugated anti-mouse secondary antibody, and nuclei were counterstained with DAPI. Coverslips were mounted with VECTASHEILD antifade mounting medium and imaged by confocal microscopy.

#### 6.4 Conservation Analysis of crRNA Target Sites in H5N1 Viral Genomes

To assess the sequence conservation of crRNA target sites within the nucleoprotein (NP) and polymerase basic protein 1 (PB1) segments of H5N1 influenza A virus, a retrospective analysis was performed using publicly available viral genomes from the GISAID database. Independent searches for NP and PB1 sequences were conducted using the following filters: A/H5N1 subtype, original passage history, highly pathogenic avian influenza (HPAI) designation, and complete genome status. Retrieved sequences were aligned to the spacer regions of crRNA_M4 (PB1-targeting) and crRNA_M10 (NP-targeting) to identify nucleotide mismatches across diverse H5N1 strains. Conservation patterns and sequence divergence were visualised using phylogenetic trees, enabling evaluation of crRNA compatibility with circulating viral variants.

### 7. Data Analysis

Data from target-specific and collateral activity screens were analysed using two-way and one-way ANOVA in GraphPad Prism (latest version), with comparisons made against GFP-only and non-targeting (NT) crRNA controls. All other assay datasets except for the data obtained from direct cDNA sequencing were evaluated using one-way ANOVA, with statistical comparisons performed relative to NT crRNA-expressing cells. For analyses involving multiplexed versus single crRNA-expressing cell lines, pairwise comparisons were conducted between each respective cell lines. The data obtained from the direct cDNA sequencing was analysed for viral RNA intermediate’s abundance (calculated as the percentage of viral-mapped reads) and compared between target specific crRNA cell lines versus NT crRNA controls using paired t-tests with Benjamini-Hochberg (BH) correction for multiple comparisons.

## Data Availability

Data obtained from flow cytometer in this study have been analysed using the latest version of FlowLogic tool. Raw files obtained from the study are available from the corresponding author upon request.

Direct cDNA sequencing data generated in this study using Oxford Nanopore Technologies (ONT) long-read sequencing have been deposited in the NCBI BioProject under accession PRJNA1392059. Raw sequencing reads are available without restriction. Processed data, including consensus sequences, transcript annotations and analysis scripts, are available from the corresponding author upon reasonable request.

Genome sequences of highly pathogenic avian influenza A(H5N1) viruses analysed in this study were obtained from the GISAID EpiFlu™ database. The sequence dataset is available under GISAID identifier EPI_SET_260107bp (DOI: https://doi.org/10.55876/gis8.260107bp). We gratefully acknowledge all data contributors via GISAID for sharing H5N1 sequence data. In accordance with GISAID data-sharing policies, raw genome sequence files are not redistributed outside the database but can be accessed directly from GISAID ^33^ subject to its terms of use. Derived data generated in this study, including multiple sequence alignments and phylogenetic tree files, are available from the corresponding author upon reasonable request.

## Acknowledgement

We thank Mrs Kirsten Morris (Senior Scientist) for her valuable technical support and for providing competent *E. coli* cells for plasmid propagation used in this study. We acknowledge Mrs Terri O’Neil for administrative support. We are grateful to Dr Jasmina Luczo and Dr Jeff Butler for providing HPAIV strains used in this work. We further thank members of the Genome Engineering Team, the Animal Biosecurity group, and all CSIRO staff at Australian Centre for Disease Preparedness Geelong for their support and assistance throughout the project. This work was financially supported by CSIRO’s internal funding through the Immune Resilience Future Science Platform.

