## Supplementary Figures for "RfxCas13d Mediates Broad-Spectrum Suppression of Highly Pathogenic Avian Influenza"

Affiliations:

**Supplementary Figures**

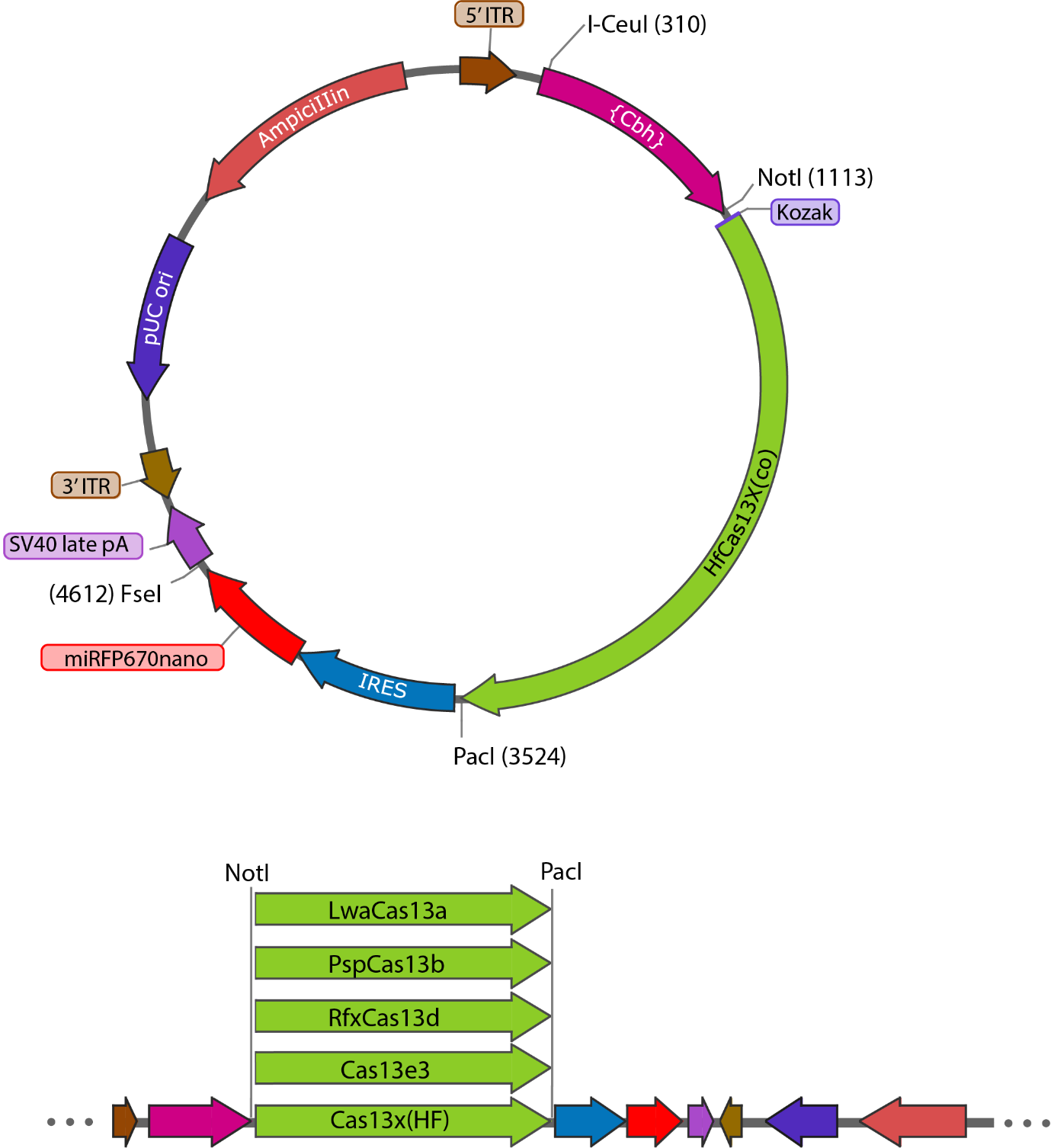

**Figure S1 | Tol2 backbone Cas13 plasmid vector map and cloning strategy.**

Schematic representation of the Tol2-based Cas13 expression vector used in this study. The map illustrates key functional elements, including the Tol2 transposon arms, Cbh promoter and regulatory sequences, restriction endonuclease sites for cloning Cas13 effector cassette, and selection markers. The cloning workflow used to insert Cas13 variant cassette into the vector is shown, assembly strategy, and orientation of inserted sequences. This figure provides an overview of the vector architecture and the steps required for constructing Cas13 expression constructs.

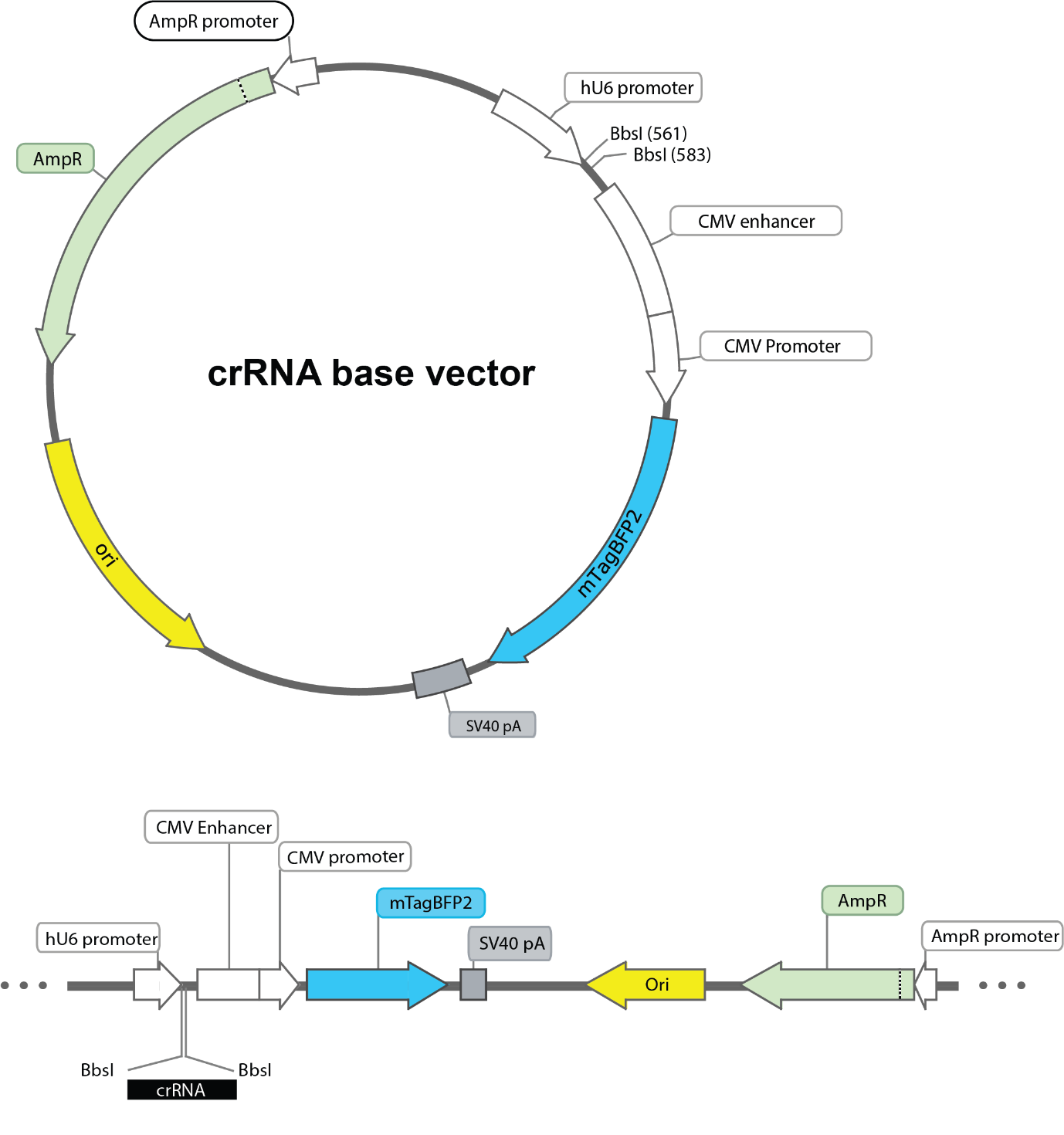

**Figure S2 | crRNA expression plasmid and cloning strategy.**

Schematic of the crRNA expression vector used in this study. The plasmid architecture includes the human U6 promoter driving crRNA expression, a *Bbs*I cloning site for guide insertion, and a CMV promoter–mTagBFP2 cassette for fluorescent selection, assembled on an ampicillin-resistant backbone.

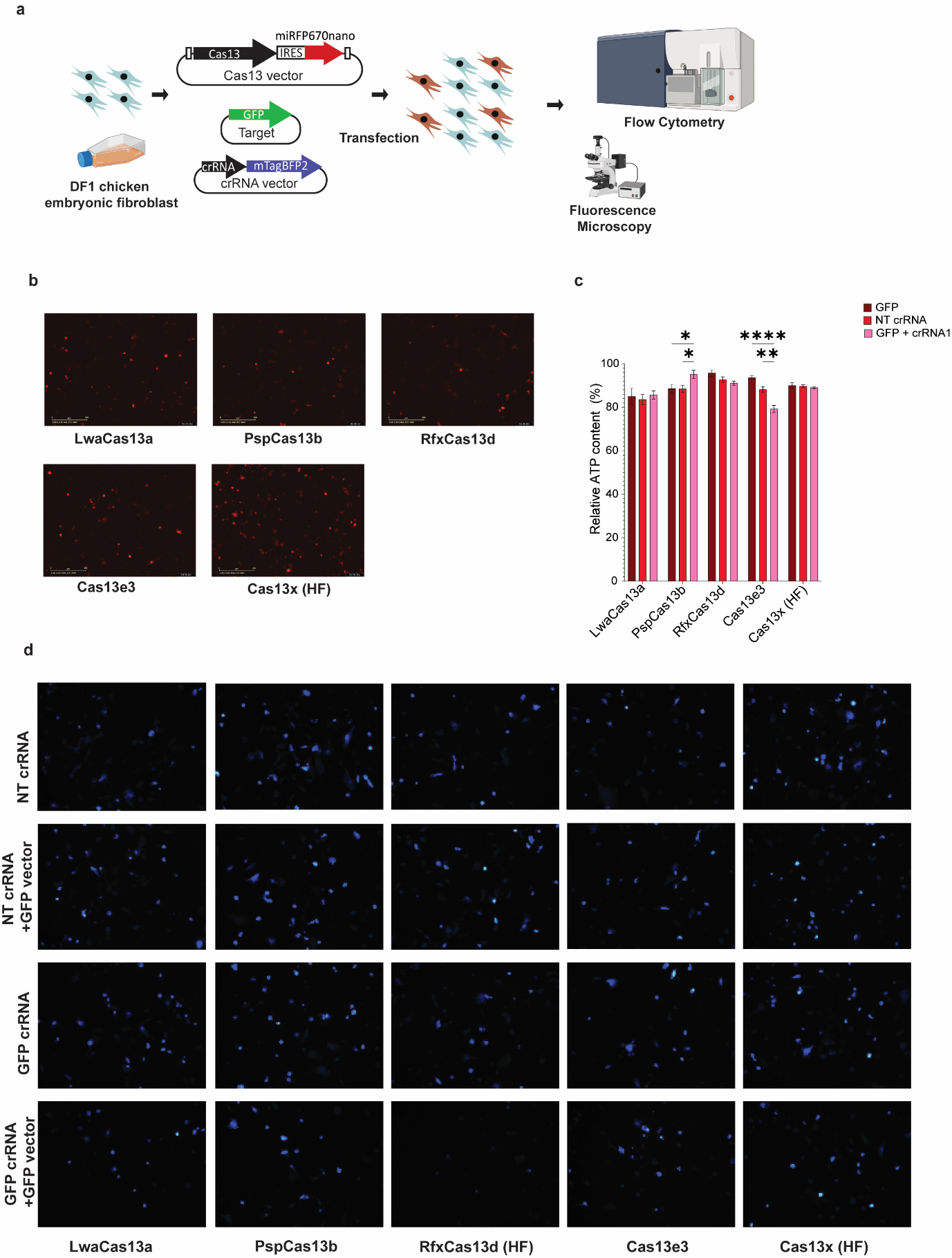

**Figure S3 | Comparative Evaluation of Cas13 Variant Expression, Target-Specific Knockdown, and Collateral RNA Cleavage in DF1 Cells**

**a** Schematic of the experimental workflow involving transient co-transfection of DF1 cells with Cas13 expression vectors (miRFP670nano, red fluorescence), crRNA vectors (mTagBFP2, blue fluorescence), and a GFP reporter plasmid (green fluorescence). Red, green, and blue fluorescence signals were used to assess Cas13 expression, target-specific knockdown, and collateral activity, respectively. **b** Representative red fluorescence images of DF1 cells transfected with plasmids encoding LwaCas13a, PspCas13b, RfxCas13d, Cas13e3 or Cas13x(HF), each co-expressing the near-infrared reporter miRFP670nano. Fluorescence intensity reflects qualitative differences in transfection efficiency and expression levels across Cas13 variants. **c** Blue fluorescence micrographs of DF1 wild-type cells co-transfected with three plasmids: a Cas13 expression vector (LwaCas13a, PspCas13b, RfxCas13d, Cas13e3 or Cas13x(HF)); a crRNA vector encoding either a target-specific crRNA (crRNA1) or a non-targeting crRNA (NT crRNA); and a GFP target vector, where indicated. Fluorescence intensity reflects collateral RNA cleavage activity triggered by each Cas13 variant in the presence or absence of target RNA. **d Relative ATP levels, measured by chemiluminescence, were used as a proxy for cell viability in DF1 cells co-transfected with Cas13 variant expression vectors and either no crRNA, a non-targeting (NT) crRNA, or a GFP-targeting crRNA. Data were analysed by two-way ANOVA using the GFP + no crRNA and GFP + NT crRNA conditions as references. Statistical significance is denoted as: *p < 0.05; **p < 0.01; ***p < 0.0001.**

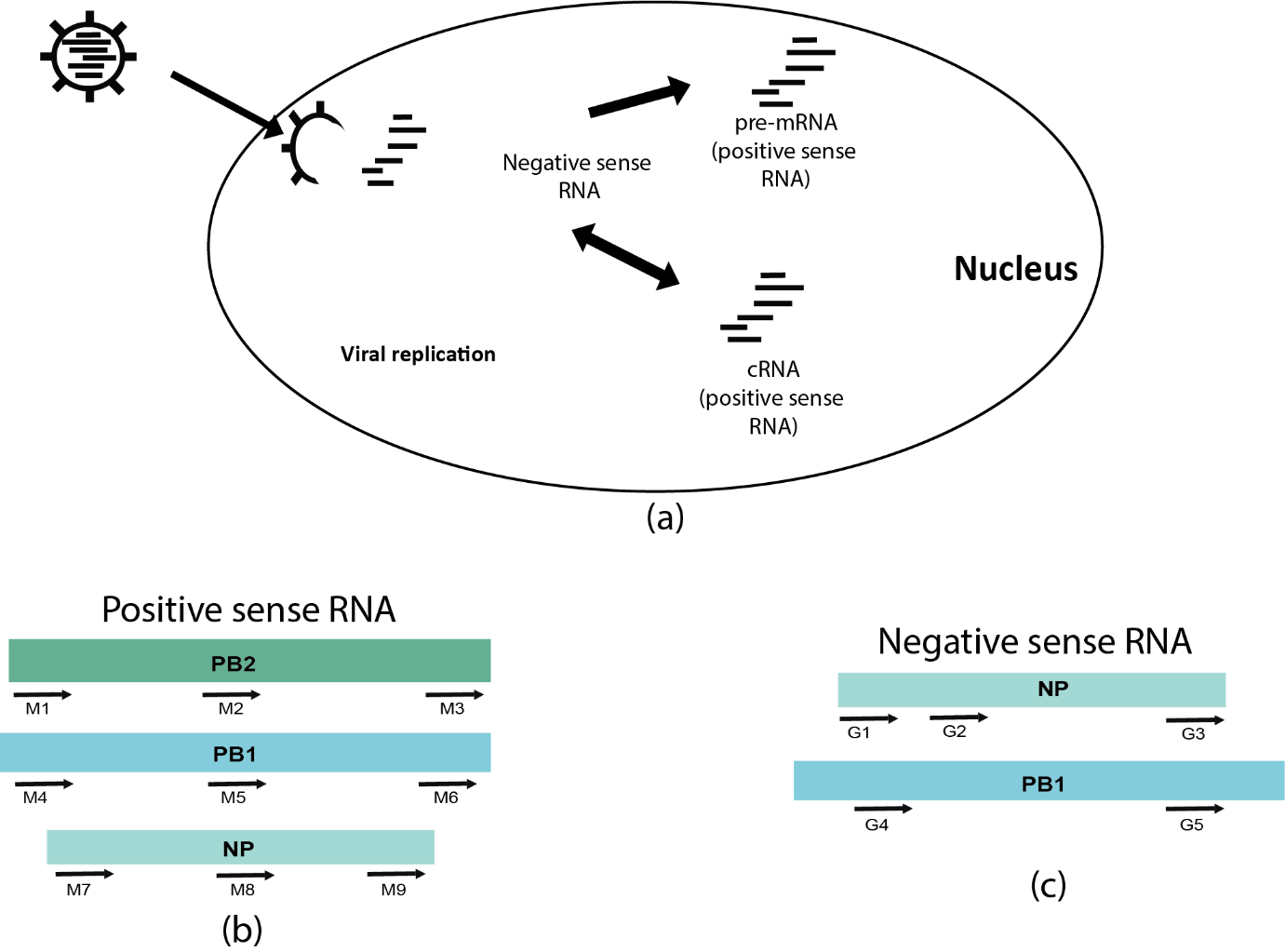

**Figure S4 | Strand-specific crRNA design strategy and target selection.**

**a, Schematic representation of influenza A virus replication within the host cell nucleus, highlighting the generation of viral RNA intermediates. b–c, Strategy for designing crRNAs targeting conserved regions of influenza viral RNA in the positive-sense (b) and negative-sense (c) strands. Positive-sense–targeting crRNAs include PB2_M1, PB2_M2, PB2_M3, PB1_M4, PB1_M5, PB1_M6, NP_M7, NP_M8, and NP_M9. Negative-sense–targeting crRNAs include NP_G1, NP_G2, NP_G3, PB1_G4, and PB1_G5.**

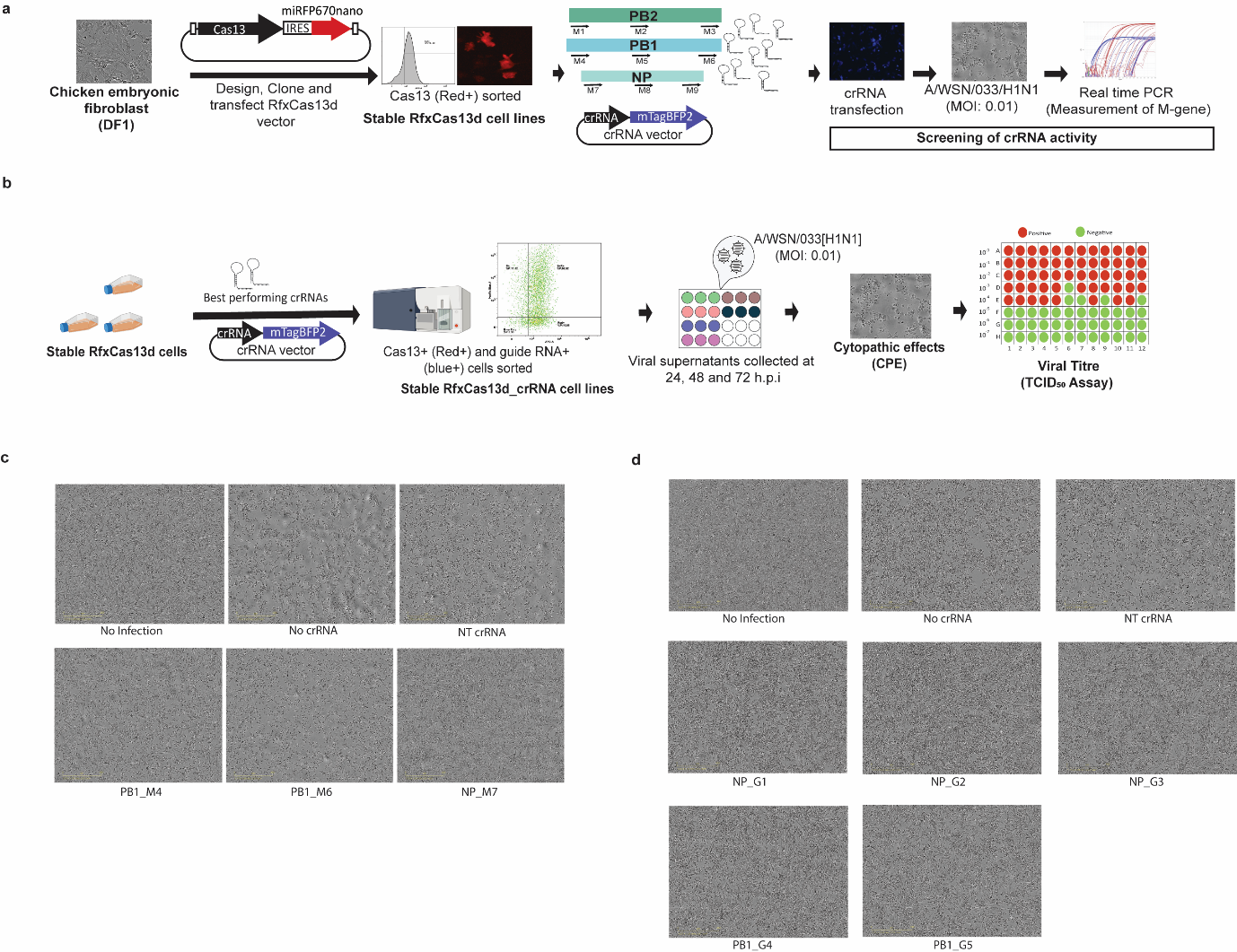

**Figure S5 | Screening and Functional Evaluation of crRNAs Targeting Influenza Viral RNA in RfxCas13d-Expressing DF1 Cells**

**a** Schematic of the screening workflow for crRNAs targeting positive sense viral RNAs. RfxCas13d cell lines were transiently transfected with crRNA vectors, followed by infection with influenza virus A/WSN/033[H1N1] at a multiplicity of infection (MOI) of 0.01. Viral M-gene expression was quantified at 24 h.p.i using TaqMan-based qRT-PCR. **b** Schematic of the generation of stable chicken DF1 cell lines co-expressing RfxCas13d and anti-influenza crRNAs. Following infection with A/WSN/033[H1N1], cytopathic effects were monitored and viral titres in supernatants were measured at defined time points. **c, d** Phase-contrast micrographs of RfxCas13d and crRNAs expressing cell lines infected with A/WSN/033 [H1N1] at an MOI of 0.01 with crRNAs targeted towards positive sense RNA at 72 h.p.i (c) and crRNAs targeted towards negative sense RNA at 24 h.p.i (d).

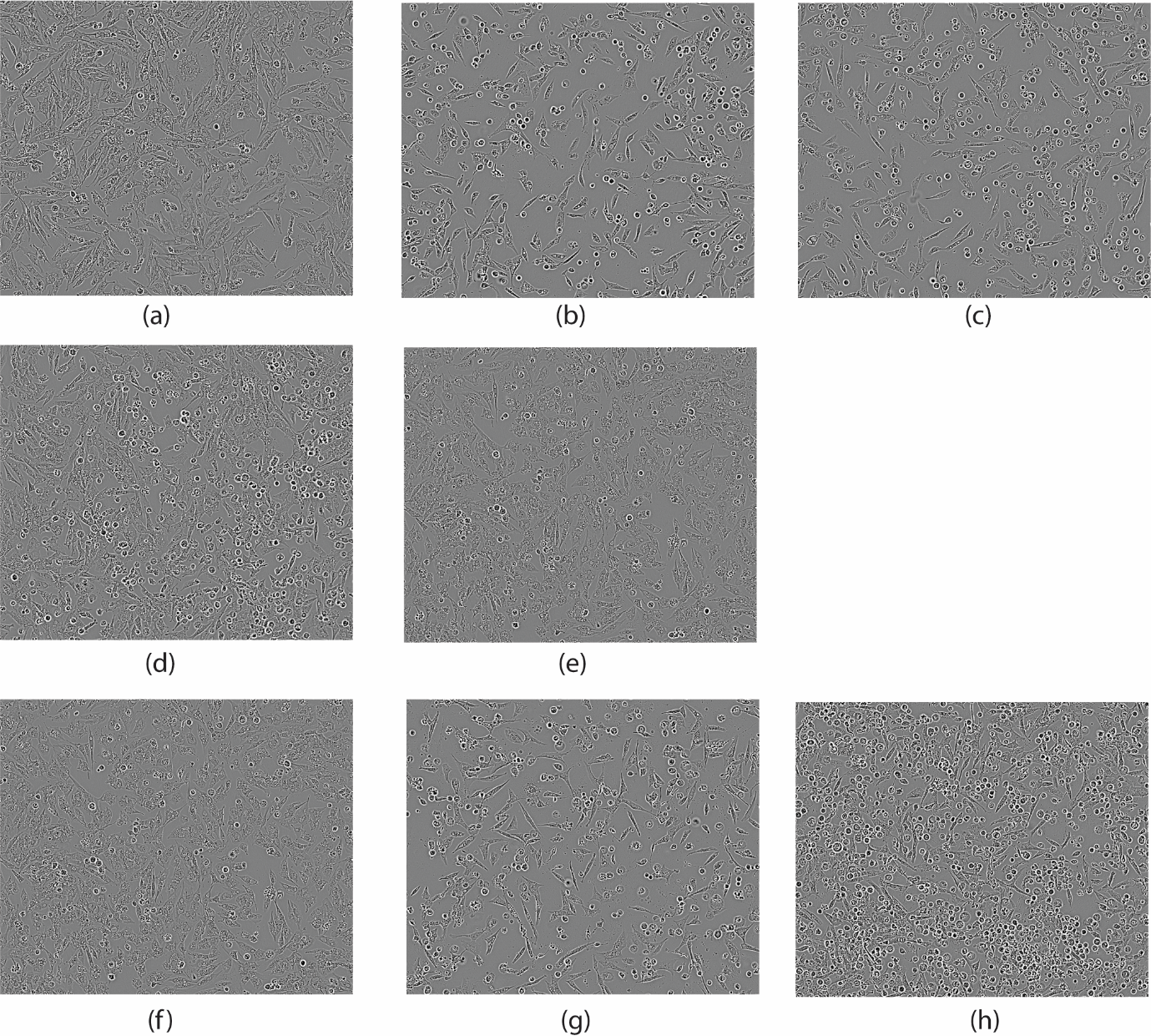

**Figure S6 | Cytopathic effects in RfxCas13d–crRNA cell lines infected with highly pathogenic H5N1 influenza virus.**

**Phase-contrast micrographs of DF1 cells at 24 h post-infection with A/Vietnam/8/2004/H5N1 (MOI = 0.01). RfxCas13d-expressing cell lines include: (a) no crRNA (uninfected control), (b) no crRNA, (c) non-targeting (NT) crRNA, and (d–h) NP_M10, PB1_M4, NP_G1, PB1_G4, and PB1_G5, respectively. Images illustrate virus-induced cytopathic effects in cells expressing guides targeting positive- and negative-sense viral RNA.**

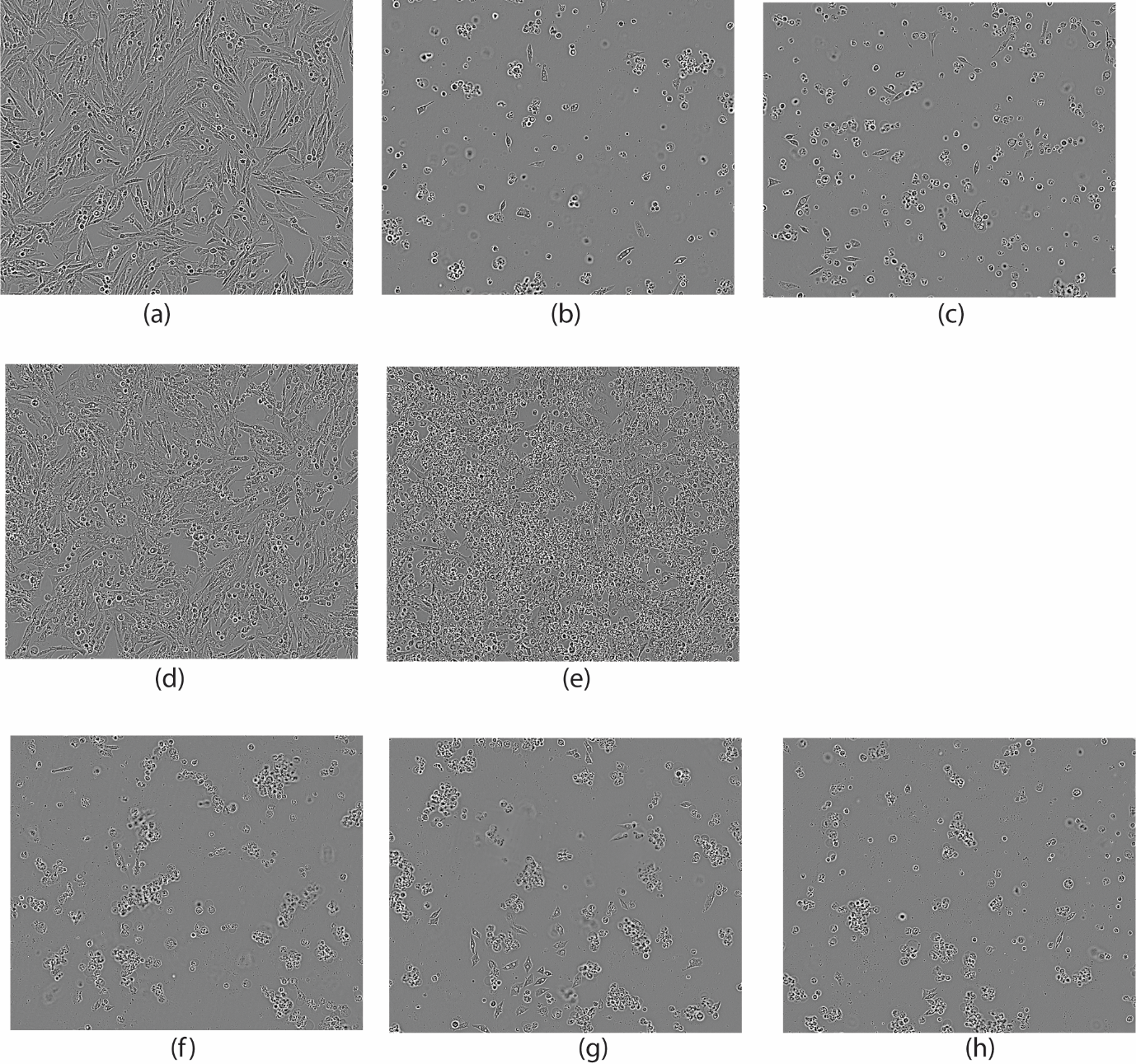

**Figure S7 | Cytopathic effects in RfxCas13d(HF)–crRNA cell lines following highly pathogenic H7N7 infection.**

**Phase-contrast micrographs of DF1 cells at 72 h post-infection with A/Lethbridge/9/2020/H7N7 (MOI = 0.01). RfxCas13d(HF)-expressing cell lines include: (a) no crRNA (uninfected control), (b) no crRNA, (c) non-targeting (NT) crRNA, and (d–h) NP_M10, PB1_M11, NP_G1, PB1_G4, and PB1_G5, respectively. Images show virus-induced cytopathic effects in cells expressing guides targeting positive- and negative-sense viral RNA.**

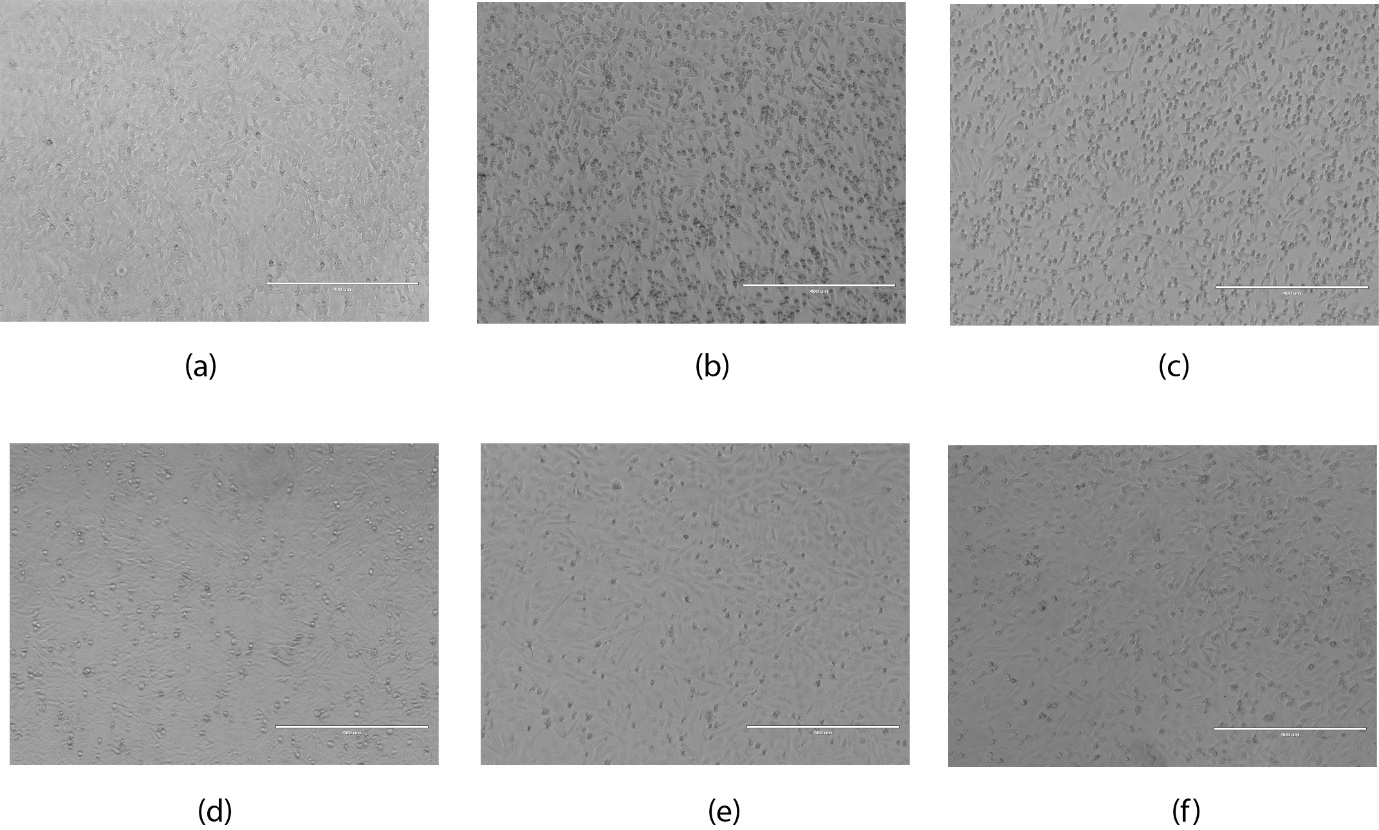

**Figure S8 | Comparison of cytopathic effects in RfxCas13d single-crRNA and crRNA-array cell lines following H5N1 infection.**

Phase-contrast micrographs of DF1 cells at 24 h.p.i with A/Chicken/Vietnam/8/2004/H5N1 (MOI = 0.01). RfxCas13d-expressing cell lines include: (a) no crRNA (uninfected control), (b) no crRNA, (c) non-targeting (NT) crRNA, and (d–f) PB1_M4, NP_M10, and PB1_NP_M4+M10, respectively. Images illustrate the relative extent of virus-induced cytopathic effects in cells expressing single or multiplexed crRNAs.

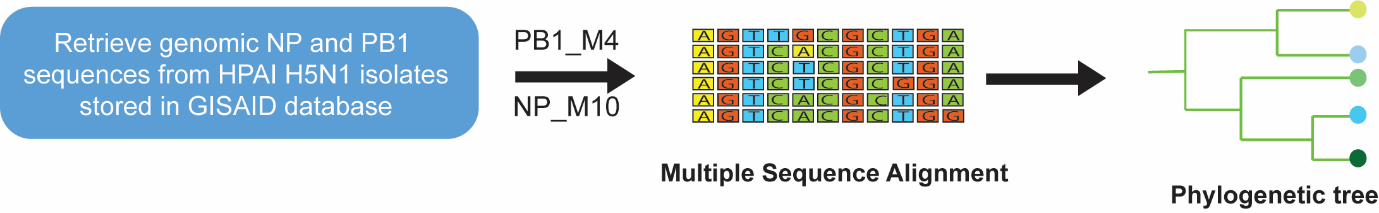

**Fig S9 | Bioinformatic pipeline testing the conservation of crRNAs across H5N1 isolates in GISAID database.**

Schematic of bioinformatic pipeline for analysing the conservation of PB1_M4 and NP_M10 crRNAs across the H5N1 isolates using multiple sequence alignment. A maximum likelihood phylogenetic tree was generated using Interactive Tree of Life (iTOL) online platform.

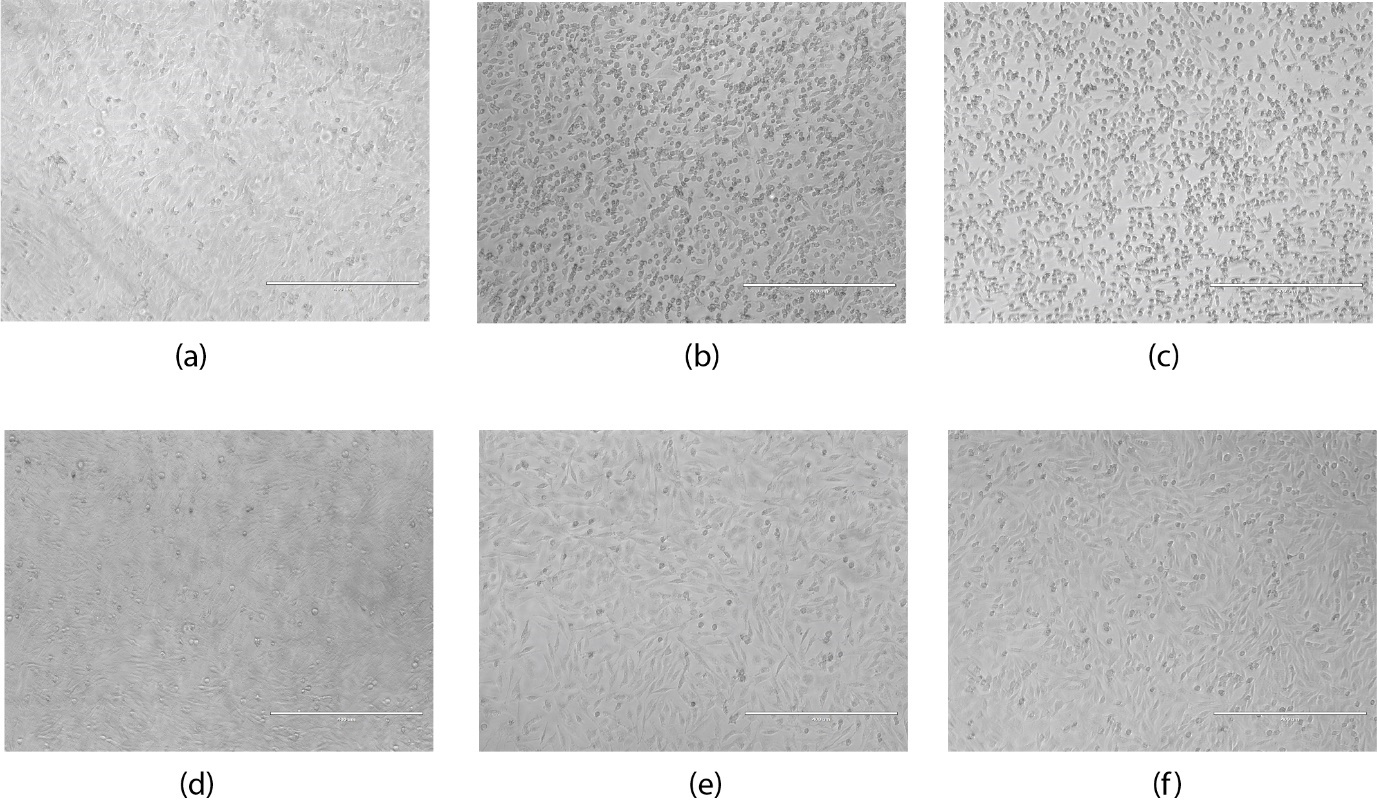

**Figure S10 | Comparison of cytopathic effects in RfxCas13d single-crRNA and crRNA-array cell lines following infection with H5N1 (a member of Clade 2.3.4.4b).**

Phase-contrast micrographs of DF1 cells at 24 h post-infection with A/Turkey/Indiana/22-003707-003/2022/H5N1 (MOI = 0.01). RfxCas13d-expressing cell lines include: (a) no crRNA (uninfected control), (b) no crRNA, (c) non-targeting (NT) crRNA, and (d–f) PB1_M4, NP_M10, and PB1_NP_M4+M10, respectively. Images depict virus-induced cytopathic effects in cells expressing single versus multiplexed crRNAs.

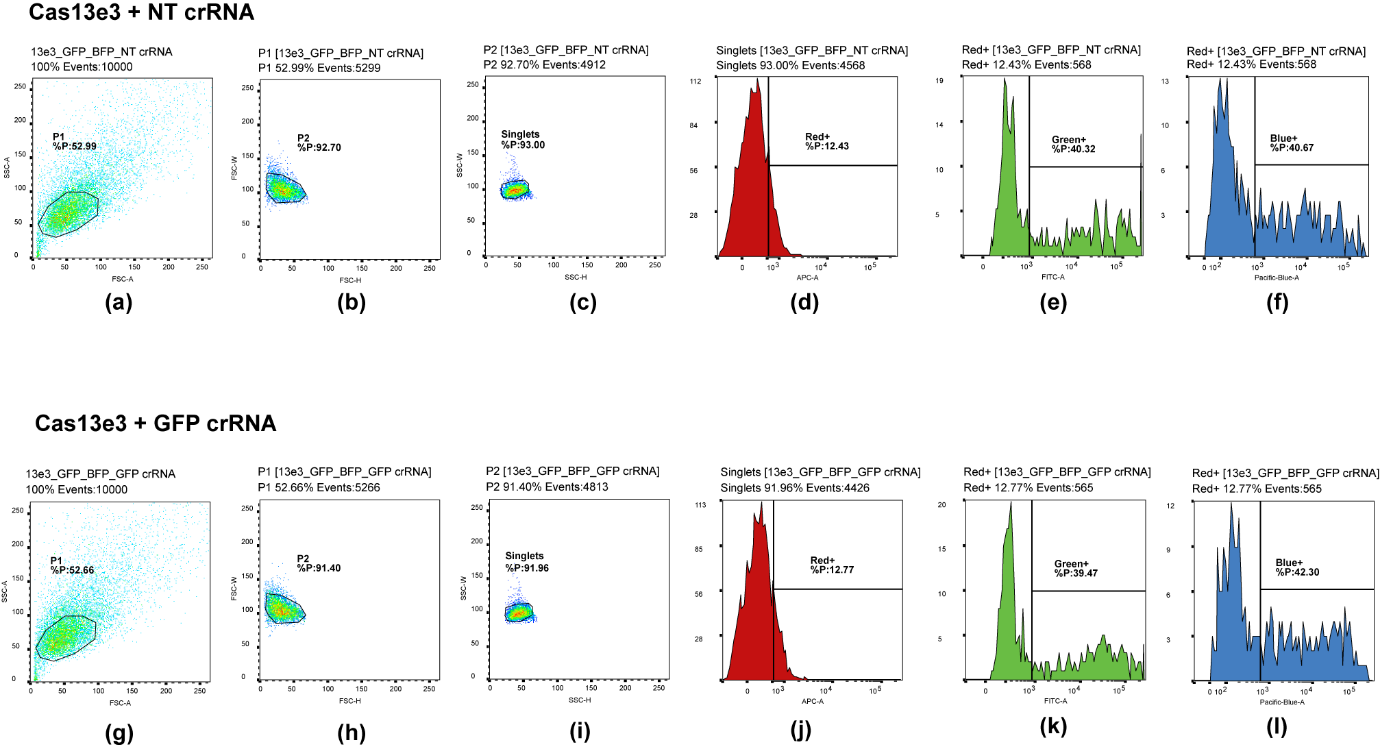

**Figure S11 | Flow cytometry gating strategy for quantifying Cas13-mediated knockdown and collateral activity.**

Schematic of the sequential gating workflow used to quantify reporter knockdown and collateral activity during Cas13 screening. Singlet cells were identified prior to measurement of green, blue and red fluorescent reporters, enabling comparative assessment of target-specific knockdown and bystander effects across Cas13 orthologs and crRNA conditions. **a–c**, Sequential gates based on forward- and side-scatter parameters—FSC-A versus SSC-A (a), FSC-H versus FSC-W (b), and SSC-H versus SSC-W (c)—were applied to exclude debris and doublets and to define singlet populations. **d–f**, Red fluorescent cells were selected in the APC-A channel (d) and subsequently analysed for green and blue reporter expression using Green⁺ (e) and Blue⁺ (f) gates for the Cas13e3 and non-targeting (NT) crRNA condition. **g–l**, Equivalent scatter (g–i) and fluorescence (j–l) gating was applied to the Cas13e3 and GFP-targeting crRNA condition, shown as a representative example of the screening workflow.

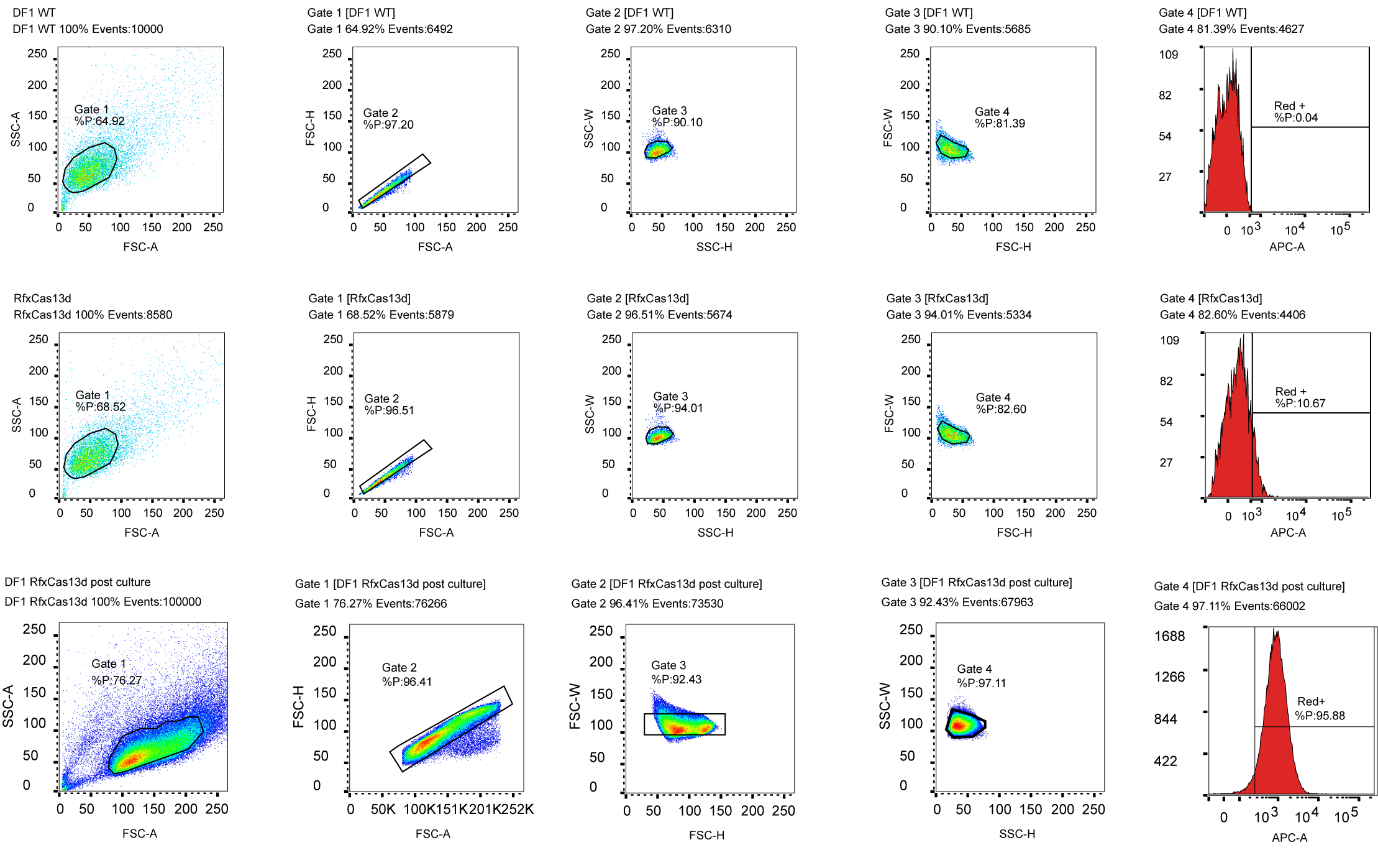

**Figure S12 | Gating strategy for fluorescence-activated cell sorting (FACS) of RfxCas13d-expressing DF1 cells and post-sort validation.**

**a–e** DF1 wild-type cells were used as negative controls to establish a sequential gating workflow comprising four scatter-based gates and a Red⁺ fluorescence gate: Gate 1, FSC-A versus SSC-A (a); Gate 2, FSC-A versus FSC-H (b); Gate 3, FSC-H versus FSC-W (c); Gate 4, SSC-H versus SSC-W (d); and the Red⁺ population defined on the APC-A histogram (e). **f–j** The identical gates defined in wild-type controls (a–e) were applied to RfxCas13d-transfected DF1 cells prior to sorting. **k–o** Post-sort and post-culture analysis of red fluorescent cells using the same gating strategy (Gates 1–4 and Red⁺), enabling direct and consistent evaluation of the sorted populations.

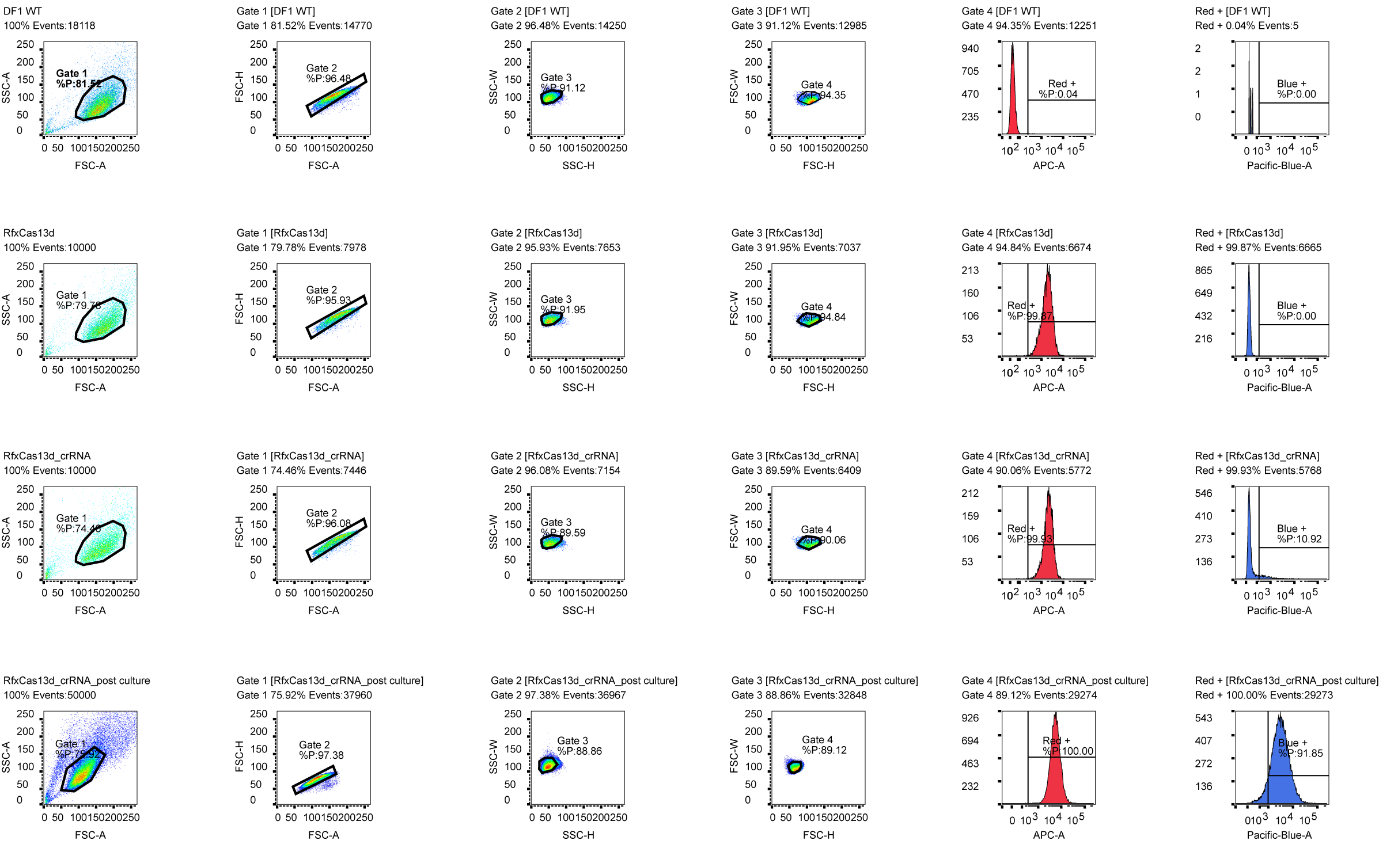

**Figure S13 | Sequential gating strategy for sorting Cas13–crRNA cell lines and post-culture validation.**

**a–f** DF1 wild-type cells were used as negative controls to establish a hierarchical gating workflow comprising four scatter-based gates and two fluorescence gates: Gate 1, FSC-A versus SSC-A (a); Gate 2, FSC-A versus FSC-H (b); Gate 3, FSC-H versus FSC-W (c); Gate 4, SSC-H versus SSC-W (d); the Red⁺ population defined on the APC-A histogram (e); and the Blue⁺ population defined on the Pacific Blue-A histogram (f).
**g–q** The identical gates defined in wild-type controls (a–f) were applied to RfxCas13d cells (g–l) and RfxCas13d crRNA-expressing cells (m–q) prior to sorting. **r–w** Post-sort and post-culture analysis of red fluorescent cells using the same gating strategy (Gates 1–4, Red⁺ and Blue⁺), enabling direct and consistent assessment of the sorted populations.
