## Supplementary Tables for "RfxCas13d Mediates Broad-Spectrum Suppression of Highly Pathogenic Avian Influenza"

Affiliations:

**Supplementary Tables**

**Table S1. Cas13 ortholog protein sequences evaluated in this study**

| **Name** | **Protein sequence** |
| --- | --- |
| LwaCas13a | MKVTKVDGISHKKYIEEGKLVKSTSENRTSERLSELLSIRLDIYIKNPDNASEEENRIRRENLKKFFSNKVLHLKDSVLYLKNRKEKNAVQDKNYSEEDISEYDLKNKNSFSVLKKILLNEDVNSEELEIFRKDVEAKLNKINSLKYSFEENKANYQKINENNVEKVGGKSKRNIIYDYYRESAKRNDYINNVQEAFDKLYKKEDIEKLFFLIENSKKHEKYKIREYYHKIIGRKNDKENFAKIIYEEIQNVNNIKELIEKIPDMSELKKSQVFYKYYLDKEELNDKNIKYAFCHFVEIEMSQLLKNYVYKRLSNISNDKIKRIFEYQNLKKLIENKLLNKLDTYVRNCGKYNYYLQVGEIATSDFIARNRQNEAFLRNIIGVSSVAYFSLRNILETENENDITGRMRGKTVKNNKGEEKYVSGEVDKIYNENKQNEVKENLKMFYSYDFNMDNKNEIEDFFANIDEAISSIRHGIVHFNLELEGKDIFAFKNIAPSEISKKMFQNEINEKKLKLKIFKQLNSANVFNYYEKDVIIKYLKNTKFNFVNKNIPFVPSFTKLYNKIEDLRNTLKFFWSVPKDKEEKDAQIYLLKNIYYGEFLNKFVKNSKVFFKITNEVIKINKQRNQKTGHYKYQKFENIEKTVPVEYLAIIQSREMINNQDKEEKNTYIDFIQQIFLKGFIDYLNKNNLKYIESNNNNDNNDIFSKIKIKKDNKEKYDKILKNYEKHNRNKEIPHEINEFVREIKLGKILKYTENLNMFYLILKLLNHKELTNLKGSLEKYQSANKEETFSDELELINLLNLDNNRVTEDFELEANEIGKFLDFNENKIKDRKELKKFDTNKIYFDGENIIKHRAFYNIKKYGMLNLLEKIADKAKYKISLKELKEYSNKKNEIEKNYTMQQNLHRKYARPKKDEKFNDEDYKEYEKAIGNIQKYTHLKNKVEFNELNLLQGLLLKILHRLVGYTSIWERDLRFRLKGEFPENHYIEEIFNFDNSKNVKYKSGQIVEKYINFYKELYKDNVEKRSIYSDKKVKKLKQEKKDLYIRNYIAHFNYIPHAEISLLEVLENLRKLLSYDRKLKNAIMKSIVDILKEYGFVATFKIGADKKIEIQTLESEKIVHLKNLKKKKLMTDRNSEELCELVKVMFEYKALE |
| PspCas13b | MNIPALVENQKKYFGTYSVMAMLNAQTVLDHIQKVADIEGEQNENNENLWFHPVMSHLYNAKNGYDKQPEKTMFIIERLQSYFPFLKIMAENQREYSNGKYKQNRVEVNSNDIFEVLKRAFGVLKMYRDLTNHYKTYEEKLNDGCEFLTSTEQPLSGMINNYYTVALRNMNERYGYKTEDLAFIQDKRFKFVKDAYGKKKSQVNTGFFLSLQDYNGDTQKKLHLSGVGIALLICLFLDKQYINIFLSRLPIFSSYNAQSEERRIIIRSFGINSIKLPKDRIHSEKSNKSVAMDMLNEVKRCPDELFTTLSAEKQSRFRIISDDHNEVLMKRSSDRFVPLLLQYIDYGKLFDHIRFHVNMGKLRYLLKADKTCIDGQTRVRVIEQPLNGFGRLEEAETMRKQENGTFGNSGIRIRDFENMKRDDANPANYPYIVDTYTHYILENNKVEMFINDKEDSAPLLPVIEDDRYVVKTIPSCRMSTLEIPAMAFHMFLFGSKKTEKLIVDVHNRYKRLFQAMQKEEVTAENIASFGIAESDLPQKILDLISGNAHGKDVDAFIRLTVDDMLTDTERRIKRFKDDRKSIRSADNKMGKRGFKQISTGKLADFLAKDIVLFQPSVNDGENKITGLNYRIMQSAIAVYDSGDDYEAKQQFKLMFEKARLIGKGTTEPHPFLYKVFARSIPANAVEFYERYLIERKFYLTGLSNEIKKGNRVDVPFIRRDQNKWKTPAMKTLGRIYSEDLPVELPRQMFDNEIKSHLKSLPQMEGIDFNNANVTYLIAEYMKRVLDDDFQTFYQWNRNYRYMDMLKGEYDRKGSLQHCFTSVEEREGLWKERASRTERYRKQASNKIRSNRQMRNASSEEIETILDKRLSNSRNEYQKSEKVIRRYRVQDALLFLLAKKTLTELADFDGERFKLKEIMPDAEKGILSEIMPMSFTFEKGGKKYTITSEGMKLKNYGDFFVLASDKRIGNLLELVGSDIVSKEDIMEEFNKYDQCRPEISSIVFNLEKWAFDTYPELSARVDREEKVDFKSILKILLNNKNINKEQSDILRKIRNAFDHNNYPDKGVVEIKALPEIAMSIKKAFGEYAIMK |
| RfxCas13d | MIEKKKSFAKGMGVKSTLVSGSKVYMTTFAEGSDARLEKIVEGDSIRSVNEGEAFSAEMADKNAGYKIGNAKFSHPKGYAVVANNPLYTGPVQQDMLGLKETLEKRYFGESADGNDNICIQVIHNILDIEKILAEYITNAAYAVNNISGLDKDIIGFGKFSTVYTYDEFKDPEHHRAAFNNNDKLINAIKAQYDEFDNFLDNPRLGYFGQAFFSKEGRNYIINYGNECYDILALLSGLRHWVVHNNEEESRISRTWLYNLDKNLDNEYISTLNYLYDRITNELTNSFSKNSAANVNYIAETLGINPAEFAEQYFRFSIMKEQKNLGFNITKLREVMLDRKDMSEIRKNHKVFDSIRTKVYTMMDFVIYRYYIEEDAKVAAANKSLPDNEKSLSEKDIFVINLRGSFNDDQKDALYYDEANRIWRKLENIMHNIKEFRGNKTREYKKKDAPRLPRILPAGRDVSAFSKLMYALTMFLDGKEINDLLTTLINKFDNIQSFLKVMPLIGVNAKFVEEYAFFKDSAKIADELRLIKSFARMGEPIADARRAMYIDAIRILGTNLSYDELKALADTFSLDENGNKLKKGKHGMRNFIINNVISNKRFHYLIRYGDPAHLHEIAKNEAVVKFVLGRIADIQKKQGQNGKNQIDRYYETCIGKDKGKSVSEKVDALTKIITGMNYDQFDKKRSVIEDTGRENAEREKFKKIISLYLTVIYHILKNIVNINARYVIGFHCVERDAQLYKEKGYDINLKKLEEKGFSSVTKLCAGIDETAPDKRKDVEKEMAERAKESIDSLESANPKLYANYIKYSDEKKAEEFTRQINREKAKTALNAYLRNTKWNVIIREDLLRIDNKTCTLFRNKAVHLEVARYVHAYINDIAEVNSYFQLYHYIMQRIIMNERYEKSSGKVSEYFDAVNDEKKYNDRLLKLLCVPFGYCIPRFKNLSIEALFDRNEAAKFDKEKKKVSGNS |
| Cas13e3 | MRGGNKSANEGKKFDFNLKQQAIFALGSNWAQKQFELTKKTQHSHRLREKLSIFQEGDIGRLEEKIENLRNYVSHGAHSGLAPLDSDEIAAFEAIVRKAVTSYLAIPREAQNIKEEKQTKIDKIRARMKKDKPLLSFSTGPLHQQNHPELMFLLAHFLTRKQLSYLIHRVYWPKEREDDKEPIQELLLFIAQPDTIIMRSKADEDARDTWISAEEEQGFAIWNYLQKRHADEVYTAPDDHYVMRQLVAFIESHQILKGAIFMRVEVRPDEKKEGAYKRVGVYEKQGSDNSLPLNIAYNTIRVSFVEDKVEGTFSLKTLIYITALFIGRVTPDKLTDFLITELKKNRDYSHRPSPAAKEGGDIKTRVEKRLCYLLRRLNKPPQNLQEQIRFICQRINFAYQQKYGQYLDQNDYKTLENLVRYYRKPDLLSWLEGNAIGQQSGIHMGQEDSKTLNQLIKASSIEQLYLDMKSHYRYGLKYVAKNYQNWPEDKVADLANIIGVRQQKTVSGNLPNAPVGVKLSWILTEFQDEIEEFKGIIHMLSAQYAPFKFEKPDKKFKKPGDAPRSEKNKRGWAHAKPFVVKQLLLNMAWYNVKEISDNDARLAGGAFVPISDISFKRSFSGCSLRMSFGKSWRQHARKSREYLQGLIDSYGEGRREFVLSKAEEENGISKSIERMEEEARQERLLFIQAILDWEKGWLKKNKSEAERLKTAKGYVEFREIAEKESLPDTIKELRNKAFHDGFLRDTKFSDCVEPIKSIYEDLKQKHI |
| Cas13x(HF) | MAQVSKQTSKKRELSIDEYQGARKWCFTIAFNKALVNRDKNDGLFVESLLRHEKYSKHDWYDEDTRALIKCSTQAANAKAEALRNYFSHYRHSPGCLTFTAEDELRTIMERAYERAIFECRRRETEVIIEFPSLFEGDRITTAGVVFFVSFFVERRVLDRLYGAVSGLKKNEGQYKLTRKALSMYCLKDSRFTKAWDKRVLLFRDILAQLGRIPAEAYEYYHGEQGDKKRANDNEGTNPKRHKDKFIEFALHYLEAQHSEICFGRRHIVREEAGAGDEHKKHRTKGKVVVDFSKKDEDQSYYISKNNVIVRIDKNAGPRSYRMGLNELKYLVLLSLQGKGDDAIAKLYRYRQHVENILDVVKVTDKDNHVFLPRFVLEQHGIGRKAFKQRIDGRVKHVRGVWEKKKAATNEMTLHEKARDILQYVNENCTRSFNPGEYNRLLVCLVGKDVENFQAGLKRLQLAERIDGRVYSIFAQTSTINEMHQVVCDQILNRLCRIGDQKLYDYVGLGKKDEIDYKQKVAWFKEHISIRRGFLRKKFWYDSKKGFAKLVEEHLESGGGQRDVGLDKKYYHIDAIGRFEGANPALYETLARDRLCLMMAQYFLGSVRKELGNKIVWSNDSIELPVEGSVGNEKSIVFSVSDYGKLYVLDDAEFLGRICEYFMPHEKGKIRAHTVAEKGFRAYNDLQKKCVEAVLAFEEKVVKAKKMSEKEGAHYIDFREILAQTMCKEAEKTAVNKVRRAFFHHHLKFVIDEFGLFSDVMKKYGIEKEWKFPVK |

Table 2: Cloning base vector sequences for Cas13 and crRNA

**Table S2. Nucleotide Sequence of Cas13 and crRNA backbone plasmid vectors**

| Plasmid Vector | Nucleotide Sequence |
| --- | --- |
| Chicken Codon optimized Cas13 base vector with Cas13x(HF) | CAGAGGTGTAAAGTACTTGAGTAATTTTACTTGATTACTGTACTTAAGTATTATTTTTGGGGATTTTTACTTTACTTGAGTACAATTAAAAATCAATACTTTTACTTTTACTTAATTACATTTTTTTAGAAAAAAAAGTACTTTTTACTCCTTACAATTTTATTTACAGTCAAAAAGTACTTATTTTTTGGAGATCACTTCATTCTATTTTCCCTTGCTATTACCAAACCAATTGAATTGCGCTGATGCCCAGTTTAATTTAAATAGATCTCAACTTTGTATAGAAAAGTTGTAACTATAACGGTCCTAAGGTAGCGACGTTACATAACTTACGGTAAATGGCCCGCCTGGCTGACCGCCCAACGACCCCCGCCCATTGACGTCAATAGTAACGCCAATAGGGACTTTCCATTGACGTCAATGGGTGGAGTATTTACGGTAAACTGCCCACTTGGCAGTACATCAAGTGTATCATATGCCAAGTACGCCCCCTATTGACGTCAATGACGGTAAATGGCCCGCCTGGCATTGTGCCCAGTACATGACCTTATGGGACTTTCCTACTTGGCAGTACATCTACGTATTAGTCATCGCTATTACCATGGTCGAGGTGAGCCCCACGTTCTGCTTCACTCTCCCCATCTCCCCCCCCTCCCCACCCCCAATTTTGTATTTATTTATTTTTTAATTATTTTGTGCAGCGATGGGGGCGGGGGGGGGGGGGGGGCGCGCGCCAGGCGGGGCGGGGCGGGGCGAGGGGCGGGGCGGGGCGAGGCGGAGAGGTGCGGCGGCAGCCAATCAGAGCGGCGCGCTCCGAAAGTTTCCTTTTATGGCGAGGCGGCGGCGGCGGCGGCCCTATAAAAAGCGAAGCGCGCGGCGGGCGGGAGTCGCTGCGCGCTGCCTTCGCCCCGTGCCCCGCTCCGCCGCCGCCTCGCGCCGCCCGCCCCGGCTCTGACTGACCGCGTTACTCCCACAGGTGAGCGGGCGGGACGGCCCTTCTCCTCCGGGCTGTAATTAGCTGAGCAAGAGGTAAGGGTTTAAGGGATGGTTGGTTGGTGGGGTATTAATGTTTAATTACCTGGAGCACCTGCCTGAAATCACTTTTTTTCAGGCGGCCGCCAAGTTTGTACAAAAAAGCAGGCTGCCACCATGCCGAAGAAAAAAAGAAAAGTGGCGCAGGTGTCTAAGCAAACATCAAAAAAACGCGAGCTGTCTATAGACGAATACCAGGGTGCACGGAAGTGGTGCTTTACCATCGCTTTCAATAAAGCACTTGTTAACCGGGACAAAAACGATGGTTTGTTTGTGGAAAGCCTCCTTCGCCATGAGAAATACTCAAAGCACGACTGGTATGACGAGGATACTCGGGCGCTCATCAAATGCTCAACGCAAGCCGCAAATGCCAAGGCAGAAGCATTGAGAAATTACTTCAGCCACTACCGCCACTCCCCTGGTTGCCTGACTTTCACGGCCGAGGACGAACTCCGGACGATAATGGAAAGAGCGTATGAGCGCGCGATATTCGAATGCAGGCGGAGAGAGACAGAGGTTATTATTGAATTCCCCAGCCTCTTCGAGGGTGATCGGATAACAACTGCAGGCGTGGTTTTCTTCGTGAGCTTTTTTGTGGAAAGGAGGGTGCTCGATCGGCTCTACGGCGCGGTTAGTGGTTTGAAAAAGAATGAAGGACAATATAAACTGACGAGGAAAGCTCTCTCCATGTACTGTCTGAAGGATAGTAGATTTACCAAAGCATGGGACAAACGCGTGCTTCTCTTTCGCGATATATTGGCTCAGTTGGGCAGGATCCCAGCTGAGGCCTATGAATATTATCACGGGGAACAGGGAGACAAGAAGCGGGCCAACGATAACGAAGGTACGAATCCAAAGAGACATAAGGATAAATTTATTGAATTTGCACTTCACTACCTTGAGGCTCAACATTCTGAAATCTGCTTCGGCCGCCGCCACATAGTACGCGAGGAGGCCGGGGCGGGTGATGAGCATAAGAAACACCGGACTAAAGGCAAGGTCGTTGTCGACTTTTCCAAAAAGGACGAGGATCAAAGCTATTACATAAGTAAGAATAATGTCATAGTCAGAATAGACAAAAATGCCGGACCGAGGTCTTACCGGATGGGGTTGAACGAGCTCAAATACTTGGTGCTCTTGTCATTGCAGGGTAAGGGAGATGACGCTATCGCCAAGCTTTACAGATATAGACAACATGTCGAAAATATCCTCGATGTAGTCAAAGTAACGGACAAGGATAACCACGTGTTCTTGCCTAGATTCGTCCTCGAACAGCACGGAATAGGCCGCAAAGCCTTTAAGCAACGGATTGACGGAAGGGTAAAACACGTTAGGGGTGTATGGGAGAAAAAAAAAGCGGCAACCAATGAAATGACGCTGCATGAAAAGGCCCGGGACATTCTGCAGTACGTAAACGAGAACTGCACTAGGAGTTTTAACCCAGGGGAATACAATAGGCTGCTGGTATGCCTCGTGGGCAAGGACGTCGAGAATTTTCAAGCTGGTCTCAAAAGATTGCAGTTGGCTGAAAGGATAGACGGACGCGTATATAGCATCTTCGCACAGACCAGCACCATTAACGAAATGCATCAGGTCGTATGCGATCAAATCCTTAACAGGCTGTGCCGGATCGGCGACCAAAAATTGTATGACTACGTTGGACTTGGCAAAAAGGACGAGATTGATTACAAGCAGAAAGTTGCATGGTTTAAAGAACATATATCCATTCGCCGGGGTTTCCTGAGGAAGAAATTCTGGTATGACAGCAAAAAGGGGTTCGCAAAATTGGTGGAGGAACACCTTGAATCTGGTGGCGGTCAGCGCGATGTCGGATTGGATAAGAAGTACTACCATATCGATGCGATCGGAAGGTTTGAGGGAGCCAACCCTGCTCTCTATGAAACTCTGGCGAGGGATCGCTTGTGCCTGATGATGGCTCAGTACTTCCTCGGATCCGTCCGCAAGGAATTGGGCAATAAAATCGTGTGGTCAAACGATAGTATAGAACTCCCAGTAGAGGGGTCAGTAGGTAATGAAAAAAGTATTGTATTTTCCGTGAGTGATTACGGTAAGCTTTACGTCCTGGACGACGCAGAATTTCTCGGACGCATATGTGAGTATTTTATGCCACATGAAAAGGGAAAGATAAGAGCTCATACCGTGGCTGAGAAAGGTTTTAGGGCGTACAACGATCTCCAGAAGAAGTGCGTAGAAGCCGTACTGGCCTTTGAGGAGAAAGTCGTGAAGGCTAAGAAGATGTCTGAGAAGGAGGGTGCCCACTATATCGATTTTAGGGAGATACTGGCCCAAACCATGTGCAAAGAAGCGGAAAAGACAGCCGTCAACAAGGTGCGGCGGGCTTTTTTTCACCACCATCTTAAGTTTGTGATAGACGAATTCGGTCTCTTCTCAGATGTGATGAAGAAATACGGCATAGAAAAAGAATGGAAATTCCCGGTTAAACCGAAGAAAAAAAGAAAAGTGTAATTAATTAAACCCAGCTTTCTTGTACAAAGTGGGCCCCTCTCCCTCCCCCCCCCCTAACGTTACTGGCCGAAGCCGCTTGGAATAAGGCCGGTGTGCGTTTGTCTATATGTTATTTTCCACCATATTGCCGTCTTTTGGCAATGTGAGGGCCCGGAAACCTGGCCCTGTCTTCTTGACGAGCATTCCTAGGGGTCTTTCCCCTCTCGCCAAAGGAATGCAAGGTCTGTTGAATGTCGTGAAGGAAGCAGTTCCTCTGGAAGCTTCTTGAAGACAAACAACGTCTGTAGCGACCCTTTGCAGGCAGCGGAACCCCCCACCTGGCGACAGGTGCCTCTGCGGCCAAAAGCCACGTGTATAAGATACACCTGCAAAGGCGGCACAACCCCAGTGCCACGTTGTGAGTTGGATAGTTGTGGAAAGAGTCAAATGGCTCTCCTCAAGCGTATTCAACAAGGGGCTGAAGGATGCCCAGAAGGTACCCCATTGTATGGGATCTGATCTGGGGCCTCGGTGCACATGCTTTACATGTGTTTAGTCGAGGTTAAAAAAACGTCTAGGCCCCCCGAACCACGGGGACGTGGTTTTCCTTTGAAAAACACGATGATAATATGGCCACAACCATGGCGAACCTTGACAAGATGCTGAATACGACGGTGACAGAGGTACGGCAATTCCTTCAGGTGGACAGAGTCTGTGTCTTCCAGTTCGAGGAGGACTATTCTGGAGTCGTCGTAGTGGAGGCTGTTGACGATCGGTGGATAAGCATACTCAAGACTCAAGTGAGAGACCGCTATTTTATGGAAACGCGCGGAGAAGAGTATAGTCATGGAAGATACCAAGCAATAGCAGACATATATACCGCAAACTTGACCGAGTGTTATAGGGATCTCTTGACTCAGTTTCAAGTTCGGGCTATATTGGCTGTACCCATCCTTCAGGGGAAGAAGTTGTGGGGTCTGCTTGTTGCTCACCAACTTGCGGCTCCCCGCCAATGGCAAACGTGGGAAATCGACTTCCTGAAACAACAAGCCGTCGTCGTCGGCATAGCCATTCAGCAATCCTAACAACTTTATTATACATAGTTGATGGCCGGCCGCTTCGAGCAGACATGATAAGATACATTGATGAGTTTGGACAAACCACAACTAGAATGCAGTGAAAAAAATGCTTTATTTGTGAAATTTGTGATGCTATTGCTTTATTTGTAACCATTATAAGCTGCAATAAACAAGTTAACAACAACAATTGCATTCATTTTATGTTTCAGGTTCAGGGGGAGGTGTGGGAGGTTTTTTAAAGCAAGTAAAACCTCTACAAATGTGGTAGATATCAAGCTTAAACAAGAATCTCTAGTTTTCTTTCTTGCTTTTACTTTTACTTCCTTAATACTCAAGTACAATTTTAATGGAGTACTTTTTTACTTTTACTCAAGTAAGATTCTAGCCAGATACTTTTACTTTTAATTGAGTAAAATTTTCCCTAAGTACTTGTACTTTCACTTGAGTAAAATTTTTGAGTACTTTTTACACCTCTGGGCGCTCTTCCGCTTCCTCGCTCACTGACTCGCTGCGCTCGGTCGTTCGGCTGCGGCGAGCGGTATCAGCTCACTCAAAGGCGGTAATACGGTTATCCACAGAATCAGGGGATAACGCAGGAAAGAACATGTGAGCAAAAGGCCAGCAAAAGGCCAGGAACCGTAAAAAGGCCGCGTTGCTGGCGTTTTTCCATAGGCTCCGCCCCCCTGACGAGCATCACAAAAATCGACGCTCAAGTCAGAGGTGGCGAAACCCGACAGGACTATAAAGATACCAGGCGTTTCCCCCTGGAAGCTCCCTCGTGCGCTCTCCTGTTCCGACCCTGCCGCTTACCGGATACCTGTCCGCCTTTCTCCCTTCGGGAAGCGTGGCGCTTTCTCATAGCTCACGCTGTAGGTATCTCAGTTCGGTGTAGGTCGTTCGCTCCAAGCTGGGCTGTGTGCACGAACCCCCCGTTCAGCCCGACCGCTGCGCCTTATCCGGTAACTATCGTCTTGAGTCCAACCCGGTAAGACACGACTTATCGCCACTGGCAGCAGCCACTGGTAACAGGATTAGCAGAGCGAGGTATGTAGGCGGTGCTACAGAGTTCTTGAAGTGGTGGCCTAACTACGGCTACACTAGAAGAACAGTATTTGGTATCTGCGCTCTGCTGAAGCCAGTTACCTTCGGAAAAAGAGTTGGTAGCTCTTGATCCGGCAAACAAACCACCGCTGGTAGCGGTGGTTTTTTTGTTTGCAAGCAGCAGATTACGCGCAGAAAAAAAGGATCTCAAGAAGATCCTTTGATCTTTTCTACGGGGTCTGACGCTCAGTGGAACGAAAACTCACGTTAAGGGATTTTGGTCATGAGATTATCAAAAAGGATCTTCACCTAGATCCTTTTAAATTAAAAATGAAGTTTTAAATCAATCTAAAGTATATATGAGTAAACTTGGTCTGACAGTTACCAATGCTTAATCAGTGAGGCACCTATCTCAGCGATCTGTCTATTTCGTTCATCCATAGTTGCCTGACTCCCCGTCGTGTAGATAACTACGATACGGGAGGGCTTACCATCTGGCCCCAGTGCTGCAATGATACCGCGAGATCCACGCTCACCGGCTCCAGATTTATCAGCAATAAACCAGCCAGCCGGAAGGGCCGAGCGCAGAAGTGGTCCTGCAACTTTATCCGCCTCCATCCAGTCTATTAATTGTTGCCGGGAAGCTAGAGTAAGTAGTTCGCCAGTTAATAGTTTGCGCAACGTTGTTGCCATTGCTACAGGCATCGTGGTGTCACGCTCGTCGTTTGGTATGGCTTCATTCAGCTCCGGTTCCCAACGATCAAGGCGAGTTACATGATCCCCCATGTTGTGCAAAAAAGCGGTTAGCTCCTTCGGTCCTCCGATCGTTGTCAGAAGTAAGTTGGCCGCAGTGTTATCACTCATGGTTATGGCAGCACTGCATAATTCTCTTACTGTCATGCCATCCGTAAGATGCTTTTCTGTGACTGGTGAGTACTCAACCAAGTCATTCTGAGAATAGTGTATGCGGCGACCGAGTTGCTCTTGCCCGGCGTCAATACGGGATAATACCGCGCCACATAGCAGAACTTTAAAAGTGCTCATCATTGGAAAACGTTCTTCGGGGCGAAAACTCTCAAGGATCTTACCGCTGTTGAGATCCAGTTCGATGTAACCCACTCGTGCACCCAACTGATCTTCAGCATCTTTTACTTTCACCAGCGTTTCTGGGTGAGCAAAAACAGGAAGGCAAAATGCCGCAAAAAAGGGAATAAGGGCGACACGGAAATGTTGAATACTCATACTCTTCCTTTTTCAATATTATTGAAGCATTTATCAGGGTTATTGTCTCATGAGCGGATACATATTTGAATGTATTTAGAAAAATAAACAAATAGGGGTTCCGCGCACATTTCCCCGAAAAGTGCCACCTGACGTCTAAGAAACCATTATTATCATGACATTAACCTATAAAAATAGGCGTATCACGAGGCCCTTTCGTC |
| crRNA base vector with human U6 and mTAGBFP2 marker | CAGAGGTGTAAAGTACTTGAGTAATTTTACTTGATTACTGTACTTAAGTATTATTTTTGGGGATTTTTACTTTACTTGAGTACAATTAAAAATCAATACTTTTACTTTTACTTAATTACATTTTTTTAGAAAAAAAAGTACTTTTTACTCCTTACAATTTTATTTACAGTCAAAAAGTACTTATTTTTTGGAGATCACTTCATTCTATTTTCCCTTGCTATTACCAAACCAATTGAATTGCGCTGATGCCCAGTTTAATTTAAATAGATCTCAACTTTGTATAGAAAAGTTGTAACTATAACGGTCCTAAGGTAGCGACGTTACGAGGGCCTATTTCCCATGATTCCTTCATATTTGCATATACGATACAAGGCTGTTAGAGAGATAATTGGAATTAATTTGACTGTAAACACAAAGATATTAGTACAAAATACGTGACGTAGAAAGTAATAATTTCTTGGGTAGTTTGCAGTTTTAAAATTATGTTTTAAAATGGACTATCATATGCTTACCGTAACTTGAAAGTATTTCGATTTCTTGGCTTTATATATCTTGTGGAAAGGACACGTCTTCGAGAAGACCTTTTTTTTCGTGGTCGACGAGATCTCGCACCGCGGGCCCGGGATCCACCGGATCTAGATCGTTACATAACTTACGGTAAATGGCCCGCCTGGCTGACCGCCCAACGACCCCCGCCCATTGACGTCAATAATGACGTATGTTCCCATAGTAACGCCAATAGGGACTTTCCATTGACGTCAATGGGTGGAGTATTTACGGTAAACTGCCCACTTGGCAGTACATCAAGTGTATCATATGCCAAGTACGCCCCCTATTGACGTCAATGACGGTAAATGGCCCGCCTGGCATTATGCCCAGTACATGACCTTATGGGACTTTCCTACTTGGCAGTACATCTACGTATTAGTCATCGCTATTACCATGGTGATGCGGTTTTGGCAGTACATCAATGGGCGTGGATAGCGGTTTGACTCACGGGGATTTCCAAGTCTCCACCCCATTGACGTCAATGGGAGTTTGTTTTGGCACCAAAATCAACGGGACTTTCCAAAATGTCGTAACAACTCCGCCCCATTGACGCAAATGGGCGGTAGGCGTGTACGGTGGGAGGTCTATATAAGCAGAGCTCAAGTTTGTACAAAAAAGCAGGCTGCCACCATGTCTGAACTGATAAAGGAGAATATGCATATGAAGCTCTATATGGAAGGTACAGTCGATAACCATCACTTCAAATGTACCTCTGAGGGGGAAGGCAAGCCTTACGAAGGGACCCAAACCATGCGGATAAAAGTAGTTGAGGGAGGGCCTCTTCCATTTGCCTTCGACATTCTTGCTACGTCTTTTTTGTATGGGAGCAAGACGTTCATTAACCATACTCAGGGGATTCCGGATTTCTTCAAACAAAGTTTTCCTGAAGGATTCACATGGGAGCGCGTAACGACTTATGAGGACGGTGGTGTCCTCACGGCCACACAGGACACATCCCTTCAAGACGGTTGCCTGATATATAACGTGAAGATTAGGGGAGTAAACTTCACGTCAAACGGTCCAGTAATGCAGAAAAAGACGCTTGGCTGGGAAGCGTTTACTGAAACCCTTTATCCTGCGGACGGCGGGCTTGAGGGTCGGAACGACATGGCTCTGAAACTCGTCGGTGGATCTCACCTTATCGCCAACGCAAAGACGACTTATCGGAGTAAAAAACCGGCAAAGAACCTCAAAATGCCCGGAGTTTATTATGTGGACTATAGGCTGGAGCGGATTAAAGAAGCGAATAACGAAACTTACGTCGAGCAGCATGAGGTCGCCGTCGCCAGATACTGTGACCTTCCATCCAAGCTGGGGCATAAACTCAATTGAACCCAGCTTTCTTGTACAAAGTGGTGATGGCCGGCCGCTTCGAGCAGACATGATAAGATACATTGATGAGTTTGGACAAACCACAACTAGAATGCAGTGAAAAAAATGCTTTATTTGTGAAATTTGTGATGCTATTGCTTTATTTGTAACCATTATAAGCTGCAATAAACAAGTTAACAACAACAATTGCATTCATTTTATGTTTCAGGTTCAGGGGGAGGTGTGGGAGGTTTTTTAAAGCAAGTAAAACCTCTACAAATGTGGTAGATATCAAGCTTAAACAAGAATCTCTAGTTTTCTTTCTTGCTTTTACTTTTACTTCCTTAATACTCAAGTACAATTTTAATGGAGTACTTTTTTACTTTTACTCAAGTAAGATTCTAGCCAGATACTTTTACTTTTAATTGAGTAAAATTTTCCCTAAGTACTTGTACTTTCACTTGAGTAAAATTTTTGAGTACTTTTTACACCTCTGGGCGCTCTTCCGCTTCCTCGCTCACTGACTCGCTGCGCTCGGTCGTTCGGCTGCGGCGAGCGGTATCAGCTCACTCAAAGGCGGTAATACGGTTATCCACAGAATCAGGGGATAACGCAGGAAAGAACATGTGAGCAAAAGGCCAGCAAAAGGCCAGGAACCGTAAAAAGGCCGCGTTGCTGGCGTTTTTCCATAGGCTCCGCCCCCCTGACGAGCATCACAAAAATCGACGCTCAAGTCAGAGGTGGCGAAACCCGACAGGACTATAAAGATACCAGGCGTTTCCCCCTGGAAGCTCCCTCGTGCGCTCTCCTGTTCCGACCCTGCCGCTTACCGGATACCTGTCCGCCTTTCTCCCTTCGGGAAGCGTGGCGCTTTCTCATAGCTCACGCTGTAGGTATCTCAGTTCGGTGTAGGTCGTTCGCTCCAAGCTGGGCTGTGTGCACGAACCCCCCGTTCAGCCCGACCGCTGCGCCTTATCCGGTAACTATCGTCTTGAGTCCAACCCGGTAAGACACGACTTATCGCCACTGGCAGCAGCCACTGGTAACAGGATTAGCAGAGCGAGGTATGTAGGCGGTGCTACAGAGTTCTTGAAGTGGTGGCCTAACTACGGCTACACTAGAAGAACAGTATTTGGTATCTGCGCTCTGCTGAAGCCAGTTACCTTCGGAAAAAGAGTTGGTAGCTCTTGATCCGGCAAACAAACCACCGCTGGTAGCGGTGGTTTTTTTGTTTGCAAGCAGCAGATTACGCGCAGAAAAAAAGGATCTCAAGAAGATCCTTTGATCTTTTCTACGGGGTCTGACGCTCAGTGGAACGAAAACTCACGTTAAGGGATTTTGGTCATGAGATTATCAAAAAGGATCTTCACCTAGATCCTTTTAAATTAAAAATGAAGTTTTAAATCAATCTAAAGTATATATGAGTAAACTTGGTCTGACAGTTACCAATGCTTAATCAGTGAGGCACCTATCTCAGCGATCTGTCTATTTCGTTCATCCATAGTTGCCTGACTCCCCGTCGTGTAGATAACTACGATACGGGAGGGCTTACCATCTGGCCCCAGTGCTGCAATGATACCGCGAGATCCACGCTCACCGGCTCCAGATTTATCAGCAATAAACCAGCCAGCCGGAAGGGCCGAGCGCAGAAGTGGTCCTGCAACTTTATCCGCCTCCATCCAGTCTATTAATTGTTGCCGGGAAGCTAGAGTAAGTAGTTCGCCAGTTAATAGTTTGCGCAACGTTGTTGCCATTGCTACAGGCATCGTGGTGTCACGCTCGTCGTTTGGTATGGCTTCATTCAGCTCCGGTTCCCAACGATCAAGGCGAGTTACATGATCCCCCATGTTGTGCAAAAAAGCGGTTAGCTCCTTCGGTCCTCCGATCGTTGTCAGAAGTAAGTTGGCCGCAGTGTTATCACTCATGGTTATGGCAGCACTGCATAATTCTCTTACTGTCATGCCATCCGTAAGATGCTTTTCTGTGACTGGTGAGTACTCAACCAAGTCATTCTGAGAATAGTGTATGCGGCGACCGAGTTGCTCTTGCCCGGCGTCAATACGGGATAATACCGCGCCACATAGCAGAACTTTAAAAGTGCTCATCATTGGAAAACGTTCTTCGGGGCGAAAACTCTCAAGGATCTTACCGCTGTTGAGATCCAGTTCGATGTAACCCACTCGTGCACCCAACTGATCTTCAGCATCTTTTACTTTCACCAGCGTTTCTGGGTGAGCAAAAACAGGAAGGCAAAATGCCGCAAAAAAGGGAATAAGGGCGACACGGAAATGTTGAATACTCATACTCTTCCTTTTTCAATATTATTGAAGCATTTATCAGGGTTATTGTCTCATGAGCGGATACATATTTGAATGTATTTAGAAAAATAAACAAATAGGGGTTCCGCGCACATTTCCCCGAAAAGTGCCACCTGACGTCTAAGAAACCATTATTATCATGACATTAACCTATAAAAATAGGCGTATCACGAGGCCCTTTCGTC |

**Table S3. crRNA constructs used to assess target-specific and collateral activity in fluorescence reporter assays**

| **Cas13 variant** | **Target** | **crRNA Sequence (**DR **+ Spacer)** |
| --- | --- | --- |
| LwaCas13a | GFP | GATTTAGACTACCCCAAAAACGAAGGGGACTAAAAC **GTCCTCCTTGAAGTCGATGCCCTTCAGC** |
|  | Non-targeting | GATTTAGACTACCCCAAAAACGAAGGGGACTAAAAC **CGTCTGGCCTTCCTGTAGCCAGCTTTCA** |
| PspCas13b | GFP | **GTCCTCCTTGAAGTCGATGCCCTTCAGC** AGTTGTGGAAGGTCCAGTTTTGAGGGGCTATTACAA |
|  | Non-targeting | **GCGTCTGGCCTTCCTGTAGCCAGCTTTCA** AGTTGTGGAAGGTCCAGTTTTGAGGGGCTATTACAA |
| RfxCas13d | GFP | GCAAGTAAACCCCTACCAACTGGTCGGGGTTTGAAAC **GTCCTCCTTGAAGTCGATGCCCTTCAGC** |
|  | Non-targeting | GCAAGTAAACCCCTACCAACTGGTCGGGGTTTGAAAC **CGTCTGGCCTTCCTGTAGCCAGCTTTCA** |
| Cas13e3 | GFP | **GTCCTCCTTGAAGTCGATGCCCTTCAGC** GCTGGAGACATCCCCATTTCTGTGGGTAAGCCAGAC |
|  | Non-targeting | **GCGTCTGGCCTTCCTGTAGCCAGCTTTCA** GCTGGAGACATCCCCATTTCTGTGGGTAAGCCAGAC |
| Cas13x(HF) | GFP | **GTCCTCCTTGAAGTCGATGCCCTTCAGC** GCTGGAGCAGCCCCCGATTTGTGGGGTGATTACAGC |
|  | Non-targeting | **GCGTCTGGCCTTCCTGTAGCCAGCTTTCA** GCTGGAGCAGCCCCCGATTTGTGGGGTGATTACAGC |

**Table S4. RfxCas13d crRNAs targeting conserved regions of A/WSN/033[H1N1]**

| Target viral RNA subtype | crRNA | Target gene | Spacer sequence | Mismatches |
| --- | --- | --- | --- | --- |
| Positive sense RNA | M1 | PB2 | TCCTTAGTTCTTTTATTCTTTCCA | 0 |
|  | M2 | PB2 | CCTCATGTACTCTTATTTTCAGTG | 0 |
|  | M3 | PB2 | CCGAATTCTTTTGGTCGCTGTCTGGC | 0 |
|  | M4 | PB1 | AAGAAAAGTAAAGTCGGATTGACA | 0 |
|  | M5 | PB1 | TCCCACCAGTAAGTAGTCTTGGTG | 0 |
|  | M6 | PB1 | TCTTCAATGGTGGAACAGATCTTCA | 0 |
|  | M7 | NP | ATCGTTTGGTGCCTTTGGTCGCCA | 0 |
|  | M8 | NP | CCATTGTTCTTTGTGCAGCTGTTTG | 0 |
|  | M9 | NP | CACTGTCTCCGAAGAAATAAGATCC | 0 |
| Negative sense RNA | G1 | NP | TGGCGTCTCAAGGCACCAAACGA | 4 |
|  | G2 | NP | GCACCAAACGATCTTACGAACAG | 0 |
|  | G3 | NP | AAGGCAACGAGCCCGATCGTGCC | 0 |
|  | G4 | PB1 | GTACCACATTCCCTTATACTGGAG | 3 |
|  | G5 | PB1 | GATCATGAAGATCTGTTCCACCA | 0 |
| Non-targeting | NT | - | CGTCTGGCCTTCCTGTAGCCAGCTTTCA | - |

**Table S5. RfxCas13d crRNA spacers targeting highly pathogenic avian influenza strains**

| crRNA | Adapted crRNA | Spacer sequence | Target gene | Target viruses |
| --- | --- | --- | --- | --- |
| M4 | - | AAGAAAAGTAAAGTCGGATTGACA | PB1 | A/Chicken/Vietnam/008/2004[H5N1] A/Turkey/Indiana/22-003707-003/2022[H5N1] (Clade 2.3.4.4b) |
| M10 | M7 | ATCGTTTGGTGCCTTGAGACGCCA | NP | A/Chicken/Lethbridge/9/2020[H7N7] A/Chicken/Vietnam/008/2004[H5N1] A/Turkey/Indiana/22-003707-003/2022[H5N1] (Clade 2.3.4.4b) |
| M11 | M4 | AAGAAAAGTAGAGTCGGATTGACA | PB1 | A/Chicken/Lethbridge/9/2020[H7N7] |
| G1 | - | TGGCGTCTCAAGGCACCAAACGA | NP | A/Chicken/Lethbridge/9/2020[H7N7] A/Chicken/Vietnam/008/2004[H5N1] A/Turkey/Indiana/22-003707-003/2022[H5N1] (Clade 2.3.4.4b) |
| G4 | - | GTACCACATTCCCTTATACTGGAG | PB1 | A/Chicken/Lethbridge/9/2020[H7N7] A/Chicken/Vietnam/008/2004[H5N1] A/Turkey/Indiana/22-003707-003/2022 [H5N1] (Clade 2.3.4.4b) |
| G5 | - | GATCATGAAGATCTGTTCCACCA | PB1 | A/Chicken/Lethbridge/9/2020[H7N7] A/Chicken/Vietnam/008/2004[H5N1] A/Turkey/Indiana/22-003707-003/2022[H5N1] (Clade 2.3.4.4b) |
| NT | - | CGTCTGGCCTTCCTGTAGCCAGCTTTCA | - | - |

**Table S6. Reverse transcription primers used for strand specific real-time PCR assay to determine the expression of HPAI genes**

| Name | Reverse Transcription primer sequence | | Target gene |
| --- | --- | --- | --- |
| NP vRNA | | AGCAAAAGCAGGGTAGATAATCACTC | NP |
| NP cRNA | | AGTAGAAACAAGGGTATTTTTCTTT |  |
| PB1 vRNA | | AAAGCAGGCAAACCATTTGAAT | PB1 |
| PB1 cRNA | | AGTAGAAACAAGGCATTTTTT |  |
| Anchored OligodT | | TTTTTTTTTTTTTTTTTTTV | mRNA |

**Table S7. TaqMan probes and primer sets for strand specific real-time PCR if influenza RNA intermediates**

| Name | TaqMan probe | Target gene | Primers |
| --- | --- | --- | --- |
| Pan flu probe | /56-FAM/TCAGGCCCCCTCAAAGCCGA/TAMRA/ | M gene | Forward: AGATGAGTCTTCTAACCGAGGTCG  Reverse:  TGCAAAGACATCTTCAAGTCTCTG |
| PB1_M4 | /5Cy5/TACTGGAGA/TAO/CCCTCCATACAGCCA/3IAbRQSp/ | PB1 crRNA_M4 region | Forward: TGAATGGATGTCAATCCGACTTT  Reverse:  TGGTGTATCCTGTCCCTGTT |
| NP_M10 | /56-FAM/AGATCGTTT/ZEN/GGTGCCTTGAGACGC/3IABkFQ/ | NP crRNA_M10 region | Forward: GCCCTGATCTCAGTAGCATTC  Reverse: CTCACCGAGTGACATCAACAT |
| PB1_G5 | /56-FAM/TCCACCATT/ZEN/GAAGAACTCAGACGGC/3IABkFQ/ | PB1 crRNA_G5 region | Forward: CACGAAGGACAAGCTAAATTCAC  Reverse: CGAGTCTGGAAGGATTAAGAAAGA |

**Table S8. Sequences of ribozymes and glycyl tRNA used in multiplex crRNA arrays**

| Name | Nucleotide Sequence |
| --- | --- |
| Chicken glycyl tRNA | GTGTTGGTGGTATAGTGGTGAGCATAGCTGCCTTCCAAGCAGTTGACCCGGGTTCGATTCCCGGCCAACGCA |
| Hammerhead ribozyme | CTGATGAGTCCGTGAGGACGAAACGAGTAAGCTCGTC |
| Hepatitis delta virus ribozyme | GGCCGGCATGGTCCCAGCCTCCTCGCTGGCGCCGGCTGGGCAACATGCTTCGGCATGGCGAATGGGAC |

**Table S9. Multiplex crRNA array sequences used in the study**

| crRNA Array | Sequence |
| --- | --- |
| glytRNA_GFP crRNA_P1 | GCAAGTAAACCCCTACCAACTGGTCGGGGTTTGAAACGTCCTCCTTGAAGTCGATGCCCTTCAGCGTGTTGGTGGTATAGTGGTGAGCATAGCTGCCTTCCAAGCAGTTGACCCGGGTTCGATTCCCGGCCAACGCAGCAAGTAAACCCCTACCAACTGGTCGGGGTTTGAAACCGTCTGGCCTTCCTGTAGCCAGCTTTCAGTGTTGGTGGTATAGTGGTGAGCATAGCTGCCTTCCAAGCAGTTGACCCGGGTTCGATTCCCGGCCAACGCAGCAAGTAAACCCCTACCAACTGGTCGGGGTTTGAAACCGTCTGGCCTTCCTGTAGCCAGCTTTCA |
| glytRNA_GFP crRNA_P2 | GCAAGTAAACCCCTACCAACTGGTCGGGGTTTGAAACCGTCTGGCCTTCCTGTAGCCAGCTTTCAGTGTTGGTGGTATAGTGGTGAGCATAGCTGCCTTCCAAGCAGTTGACCCGGGTTCGATTCCCGGCCAACGCAGCAAGTAAACCCCTACCAACTGGTCGGGGTTTGAAACGTCCTCCTTGAAGTCGATGCCCTTCAGCGTGTTGGTGGTATAGTGGTGAGCATAGCTGCCTTCCAAGCAGTTGACCCGGGTTCGATTCCCGGCCAACGCAGCAAGTAAACCCCTACCAACTGGTCGGGGTTTGAAACCGTCTGGCCTTCCTGTAGCCAGCTTTCA |
| glytRNA_GFP crRNA_P3 | GCAAGTAAACCCCTACCAACTGGTCGGGGTTTGAAACCGTCTGGCCTTCCTGTAGCCAGCTTTCAGTGTTGGTGGTATAGTGGTGAGCATAGCTGCCTTCCAAGCAGTTGACCCGGGTTCGATTCCCGGCCAACGCAGCAAGTAAACCCCTACCAACTGGTCGGGGTTTGAAACCGTCTGGCCTTCCTGTAGCCAGCTTTCAGTGTTGGTGGTATAGTGGTGAGCATAGCTGCCTTCCAAGCAGTTGACCCGGGTTCGATTCCCGGCCAACGCAGCAAGTAAACCCCTACCAACTGGTCGGGGTTTGAAACGTCCTCCTTGAAGTCGATGCCCTTCAGC |
| ribozyme_GFP crRNA_P1 | GCAAGTAAACCCCTACCAACTGGTCGGGGTTTGAAACGTCCTCCTTGAAGTCGATGCCCTTCAGCGGCCGGCATGGTCCCAGCCTCCTCGCTGGCGCCGGCTGGGCAACATGCTTCGGCATGGCGAATGGGACATGCATGCATCGACTTGCCTGATGAGTCCGTGAGGACGAAACGAGTAAGCTCGTCGCAAGTAAACCCCTACCAACTGGTCGGGGTTTGAAACCGTCTGGCCTTCCTGTAGCCAGCTTTCAGGCCGGCATGGTCCCAGCCTCCTCGCTGGCGCCGGCTGGGCAACATGCTTCGGCATGGCGAATGGGACATGCATGCATCGACTTGCCTGATGAGTCCGTGAGGACGAAACGAGTAAGCTCGTCGCAAGTAAACCCCTACCAACTGGTCGGGGTTTGAAACCGTCTGGCCTTCCTGTAGCCAGCTTTCA |
| ribozyme_GFP crRNA_P2 | GCAAGTAAACCCCTACCAACTGGTCGGGGTTTGAAACCGTCTGGCCTTCCTGTAGCCAGCTTTCAGGCCGGCATGGTCCCAGCCTCCTCGCTGGCGCCGGCTGGGCAACATGCTTCGGCATGGCGAATGGGACATGCATGCATCGACTTGCCTGATGAGTCCGTGAGGACGAAACGAGTAAGCTCGTCGCAAGTAAACCCCTACCAACTGGTCGGGGTTTGAAACGTCCTCCTTGAAGTCGATGCCCTTCAGCGGCCGGCATGGTCCCAGCCTCCTCGCTGGCGCCGGCTGGGCAACATGCTTCGGCATGGCGAATGGGACATGCATGCATCGACTTGCCTGATGAGTCCGTGAGGACGAAACGAGTAAGCTCGTCGCAAGTAAACCCCTACCAACTGGTCGGGGTTTGAAACCGTCTGGCCTTCCTGTAGCCAGCTTTCA |
| ribozyme_GFP crRNA_P3 | GCAAGTAAACCCCTACCAACTGGTCGGGGTTTGAAACCGTCTGGCCTTCCTGTAGCCAGCTTTCAGGCCGGCATGGTCCCAGCCTCCTCGCTGGCGCCGGCTGGGCAACATGCTTCGGCATGGCGAATGGGACATGCATGCATCGACTTGCCTGATGAGTCCGTGAGGACGAAACGAGTAAGCTCGTCGCAAGTAAACCCCTACCAACTGGTCGGGGTTTGAAACCGTCTGGCCTTCCTGTAGCCAGCTTTCAGGCCGGCATGGTCCCAGCCTCCTCGCTGGCGCCGGCTGGGCAACATGCTTCGGCATGGCGAATGGGACATGCATGCATCGACTTGCCTGATGAGTCCGTGAGGACGAAACGAGTAAGCTCGTCGCAAGTAAACCCCTACCAACTGGTCGGGGTTTGAAACGTCCTCCTTGAAGTCGATGCCCTTCAGC |
| PB1_NP_M4+M10 | GCAAGTAAACCCCTACCAACTGGTCGGGGTTTGAAACAAGAAAAGTAAAGTCGGATTGACAGGCCGGCATGGTCCCAGCCTCCTCGCTGGCGCCGGCTGGGCAACATGCTTCGGCATGGCGAATGGGACATGCATGCATCGACTTGCCTGATGAGTCCGTGAGGACGAAACGAGTAAGCTCGTCGCAAGTAAACCCCTACCAACTGGTCGGGGTTTGAAACATCGTTTGGTGCCTTGAGACGCCA |
